# Malignant tumors engender and employ cancer-specific organelles for self-protection and metastasis

**DOI:** 10.1101/2023.07.15.549159

**Authors:** Tingfang Yi, Gerhard Wagner

**Affiliations:** Cytocapsula Research Institute, 245 First Street, Cambridge, MA, 02142 USA; Centiver Ltd., 245 First Street, Cambridge, MA, 02142 USA; Department of Biological Chemistry and Molecular Pharmacology of Harvard Medical School, 240 Longwood, Boston, MA, 02115 USA

## Abstract

Cancer is a leading source of human lethality but current therapies have had limited efficacy in many cancers, which highlights an unmet need to assess the underlying mechanisms that govern cancer progression *in vivo.* Here we show that, upon malignant transformation, aggressive oncocells generate extracellular membranous compartments, cytocapsulas or cytocapsular tubes (CCTs), to enclose oncocells and engender cytocapsular oncocells *in vivo*. Cytocapsular oncocells are universally present in solid cancers and appear in hematologic cancers in the immune organs. Networks of cytocapsular tubes provide membrane-enclosed freeways for protected cancer metastasis. CCT networks interconnect cytocapsular tumors creating cytocapsular tumor network systems. Our findings suggest that cytocapsular oncocells drive membrane-encompassed cancer progression. Thus, interconnected cytocapsular oncocells, CCT networks and cytocapsular tumor network systems coordinate cancer progression in the integrated cytocapsular membrane systems.

## Introduction

Cancer ranks as a leading cause of mankind lethality(*1–4*), and almost 10 million cancer deaths and around 19.3 million new cancer cases occurred in 2020 alone worldwide (*1*). In the last decades, many aspects of cancer progression have been intensively studied. These include sub-clonal, cellular, cellular-plastic, genetic, genomic, proteomic, signaling, metabolic, and tumor microenvironmental topics(*5–7*). Several models for tumor progression have been hypothesized, which invoke diverse scales of progression-driving factors: genetic alterations, genomic aberrations (discordant inheritance, DNA macro-alterations), oncoprotein promotion, signaling pathways, cell plasticity, intercellular reactions, and microenvironments (*8–10*). Tumors’ biological, biochemical, biophysical and metabolic features are targeted for the invention of numerous types of cancer diagnosis and therapies(*11–12*). However, the current clinical cancer treatment outcomes suggest that the mechanisms underlying cancer development and progression *in vivo* have not been fully resolved (*1*, *7–13*).

Recently, we reported two new organelles dubbed cytocapsula (CC) and cytocapsular tube (CCT) (*14*). CCs and CCTs are two bi-phospholipid-layer membrane composed, facultative and extracellular organelles that were discovered in mammalian cells *in vitro* and in human cancer xenografts in mouse. CCTs provide membrane-enclosed physical freeways for cell migration in 3D matrices *in vitro* and in breast cancer HMLER(CD44^high^/CD24^low^)^FA^ subpopulation cells induced xenografts in mouse (*14*). Cytocapsular membranes enclose the cells, physically shielding them from the extracytocapsular microenvironments and serving as protective coverings against environmental stresses and attacks (*14*). Here, we asked whether CCs and CCTs play a role in human cancer development and progression *in vivo*.

## Results

### Detection and lifecycle of cytocapsular oncocells *in vitro* and in human tissues *in vivo*

Using CC/CCT culture kit and Unipick, we cultured and collected acellular SILAC labelled cytocapsulas (CCs) of Bxpc3 pancreas cancer cells, MCF-7 breast cancer cells, and SK-CO-1 colon cancer cells for CC proteome assays (**Fig.1A**). The most prominent protein calcium pump plasma membrane Ca^2+^-ATPase 2 (PMCA2) consistently appears in the plasma membranes of the enclosed cancer cells (**Fig.1B**). However, PMCA2 is at much higher abundance in the CC membranes encapsulating single or multiple Bxpc3 cancer cells (number tested, n=601), or in ecellulated CC membranes (n=514) *in vitro*. No PMCA2 signal is found in the CC/CCT culture kit matrix (**Fig.1B**). PMCA2 and γ-actin are constantly colocalized in cancer cell plasma membranes (n=601) and in completely ecellulated CC membranes (n=514) *in vitro* (**Fig. 1B**). In addition, PMCA2 consistently shows high abundance and colocalizes with γ-actin in the cytocapsular tube (CCT) membranes surrounding Bxpc3 cancer cell (n=106, **fig. S1A**). The same is found for the enlarged CC membranes surrounding tumorspheres (n=546, **fig.S1B**), and for the ecellulated enlarged CC membranes of tumorspheres (n=127, **fig. S1B**). These observations suggest that PMCA2 may be a molecular marker of CCs and CCTs *in vitro*. Next, we investigated human normal, benign tumor and cancer tissues with anti-PMCA2 antibodies. In clinical normal (n=14 patients, 1 tissue/patient, the same as below, **fig. S1C1**) or benign tumor tissues (n=126 patients, **fig. S1C2 and table S1**), there are very low PMCA2 signals and no CC/CCT, indicating that normal tissues and benign tumor tissues don’t generate CCs/CCTs, and that PMCA2 expression is tightly controlled and maintained at low abundance in human normal tissues or benign tumors (**fig. S1C, 1 and 2**). In contrast, in clinical breast (n=685 patients, **fig.S1C3**) and pancreas (n=310 patients, **fig.S1C4**) carcinoma tissues, there are many long and curved CCTs with high abundance of PMCA2 in the CCT membranes (**fig. S1C, 3 and 4**). In addition, there are no PMCA2 signals in the extracellular matrix (ECM) in normal, benign and cancer tissues (**fig. S1C**). The above observations suggest that PMCA2 may be a molecular marker of CCs/CCTs *in vitro* and *in vivo*.

**Fig. 1.**
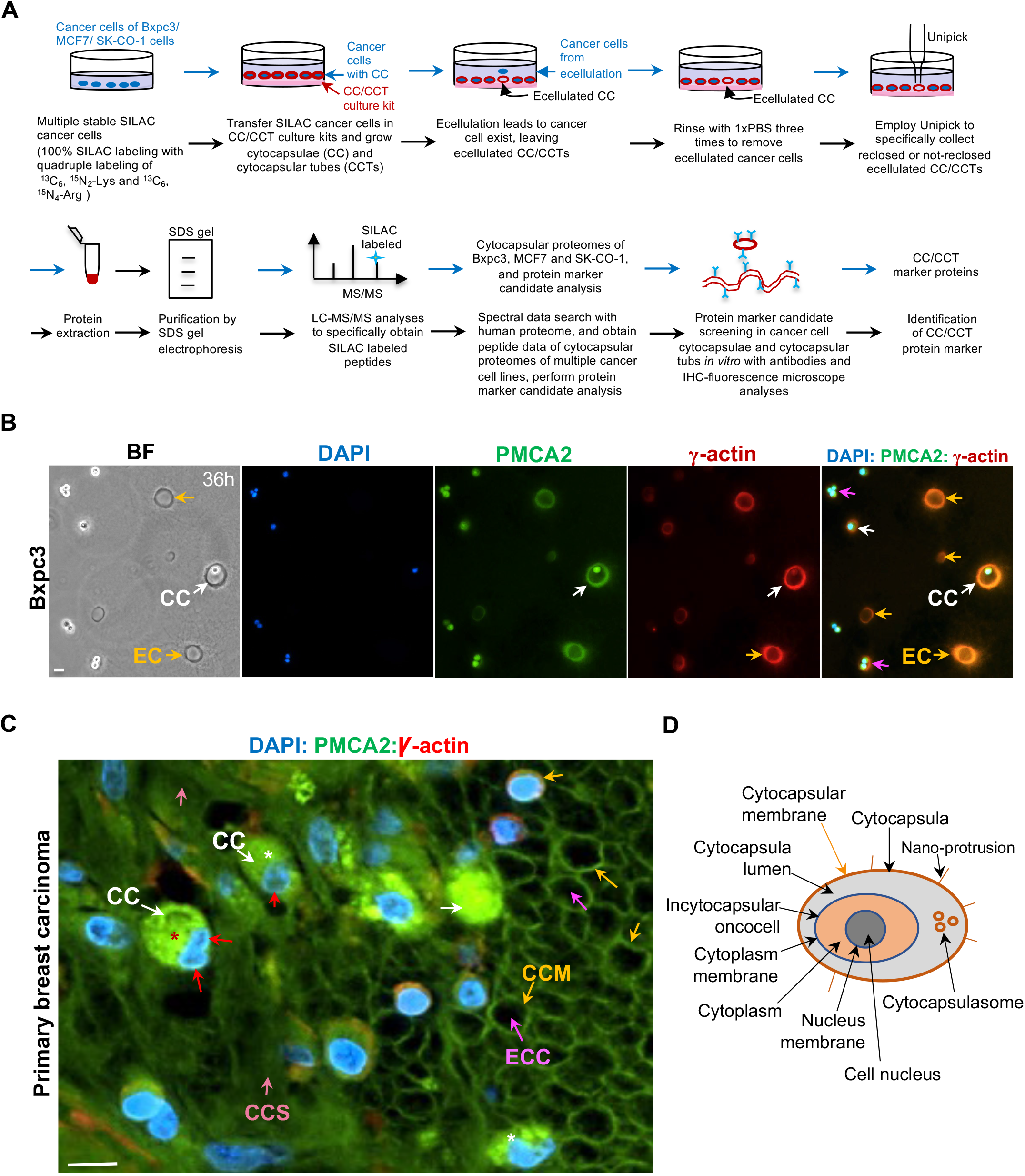
The lifecycle of cytocapsular oncocells *in vitro* and in human tissues *in vivo*. (**A**) Schematic diagram of cancer cell cytocapsular proteome analysis and cytocapsula (CC) and cytocapsular tube (CCT) molecular marker identification. (**B**) Representative immunohistochemistry (IHC) fluorescence microscope images of pancreas cancer Bxpc3 cells with cytocapsulas (CC) in the 3D CC/CCT culture kit matrix. Cancer cells with CC (white arrows), ecellulated cytocapsulas (ECC, orange arrows), and cancer cell proliferation in CC (purple arrows) were shown. (**C**) A representative IHC fluorescence microscope image of primary breast carcinoma with cytocapsular oncocells (white arrows). Incytocapsular oncocell (red arrows), ecellulated cytocapsulas (ECC, purple arrows), cytocapsular membrane (CCM, orange arrows), and cytocapsular oncocell proliferation in CC (red asterisk), and folded CC (white asterisk) are shown. (**D**) A schematic diagram of a cytocapsular oncocell with its cytocapsula, cytocapsulasome and nano-protrusions. Scale bar in B and C, 10μm.

With the PMCA2 biomarker in hand, we tested whether cancerous cells generate cytocapsulas *in vivo*. Indeed, in the early stage of breast carcinoma, there are many cancerous breast cells enclosed by CCs, and thus isolated from microenvironments (**Fig. 1C**). Some CCs (n=135) appear in highly folded shapes, with breast cancer cells wrapped inside, while others (n=268) tightly wrap breast cancer cells (**Fig.1C**). Some breast cancer cells ecellulate from the CCs (n=1023), leaving acellular CCs behind (**Fig.1C**). These acellular CCs align together and form acellular CC groups (**Fig. 1C**). Some acellular CCs appear in round or oval morphologies (n=576), and many adopt irregular shapes (n=1217) (**Fig. 1C**). Some acellular CC membranes indicate a taut and stringent status (n=107, **Fig.1C**). The CC ecellulation phenomena and acellular CCs’ morphologies *in vivo* are consistent with those *in vitro* (**Fig. 1B and fig. S1B**). Importantly, cancer cells proliferate in CC lumens and migrate in CCT lumens *in vitro* and *in vivo* (**Fig. 1, B and C, and fig. S1, A and B**). These observations suggest that the cytocapsular bi-layer phospholipid membranes not only shelter and protect cancerous cells inside but also allow cancerous cells to execute cellular behaviors and activities in the CC/CCT lumens. We termed this previously unappreciated single cancerous cell that is enclosed in an extracellular and membranous cytocapsula or cytocapsular tube (*14*), and performs cellular activities in the cytocapsular lumen, a “cytocapsular oncocell (onco-: “tumor”)” (**Fig.1D**).

Single cytocapsular oncocells generate elongated cytocapsular tubes and migrate inside (**fig. S1A**). Cytocapsular oncocells proliferate in cytocapsular lumens and grow into cytocapsular tumorspheres *in vitro* **(Fig.1B and fig.S1B)** and initiate tumor formation *in vivo* (**Fig. 1C**). Sometimes, cytocapsular oncocell ecellulation generates acytocapsular oncocells and acellular cytocapsulas followed by autodegradation into cytocapsular strands (**Fig. 1C**). The lifecycle of cytocapsular oncocells includes 3 successive procedures: 1) Incytocapsular oncocells proliferate and grow into cytocapsular tumors wrapped in enlarged cytocapsulas, 2) cytocapsulas elongate and generate cytocapsular tube with oncocells in migration inside, 3) cytocapsular ecellulation engendering acytocapsular oncocells and acellular cytocapsulas (CCs), followed by CC degradation (**fig.S1D**).

Next, we examined why CCs and CCTs were previously not recognized. In the continuously sectioned adjacent specimens from the same site of the same cancer samples, Hematoxylin and Eosin (H&E) staining does not show features for the large quantities of CCTs in cancers due to the poor staining of CCT membranes by Eosin (n=352 patients, **fig. S2**). The antibodies recognizing clinical breast cancer cell marker proteins ER, PR and HER-2 don’t recognize CCT marker proteins in these breast cancer samples (n=213patients, **fig.S3A**). Furthermore, colon cancer cell markers MSH-2 do not show features of CCTs in colon cancers, because they do not stain CCT membrane marker proteins and lack CCT membrane staining (n=86 patients, **fig. 3B**). The above observations suggest that there are 6 major possibilities why CCs and CCTs haven’t been documented for a long time: for *in vivo* assays: 1) H&E staining shows poor staining of CCs/CCTs, 2) the high-abundance CC/CCT marker proteins have previously been unknown or high quality antibodies recognizing CC/CCT marker proteins were not available, 3) the low resolution of fluorescence microscope images or a high autofluorescence noise background caused low signal/noise ratio, and CCs/CCTs were missed, 4) compact acytocapsular oncocell masses are emphasized and signals of CCs/CCTs are ignored, or treated as extracellular matrix (ECM) or unknown phenomena without further study; for *in vitro* analyses: 1) ECM mimics (such as collagen gel, Matrigel matrix) in the cell culture plates are difficult to meet the requested precisely controlled biochemical, biophysical, and biomechanical characteristics (e.g., polymerization, density, and viscoelasticity) of the ECM mimics, and 2) conventional CDX (cancer cell line derived xenografts), PDX (patient cancer cell derived xenografts) and O-PDX (orthotopic-PDX) do not generate CCs or CCTs (**fig. S2 and fig. S3, A and B**).

### Progression and lifecycle of cytocapsular tumors

We asked if cytocapsular oncocells grow into tumorspheres *in vitro*. Indeed, at 48h and 72h, in the CC/CCT culture kit matrix, cytocapsular Bxpc3 oncocells proliferate in CCs and grow into tumorspheres, and small CCs develop into enlarged CCs enclosing tumorspheres inside (n=458, **Fig. 2A and fig.S1B**). Similar like single cytocapsular oncocell ecellulation, some tumorspheres in large CCs perform spontaneous ecellulation (n=23 tumorspheres, **fig.S1B**). Individual oncocell ecellulation of tumorspheres in enlarged CCs generates large, reclosed and reunited, acellular, deflated and concave CC discs (n=35, **fig. S1B**). Ecellulation of tumorspheres in large CCs engender large, not-reclosed/reunited, acellular, deflated CCs with big open holes (n=74, **fig. S1B**). These observations suggest that solid cytocapsular oncocells grow into tumorspheres in enlarged CCs *in vitro*.

**Fig. 2.**
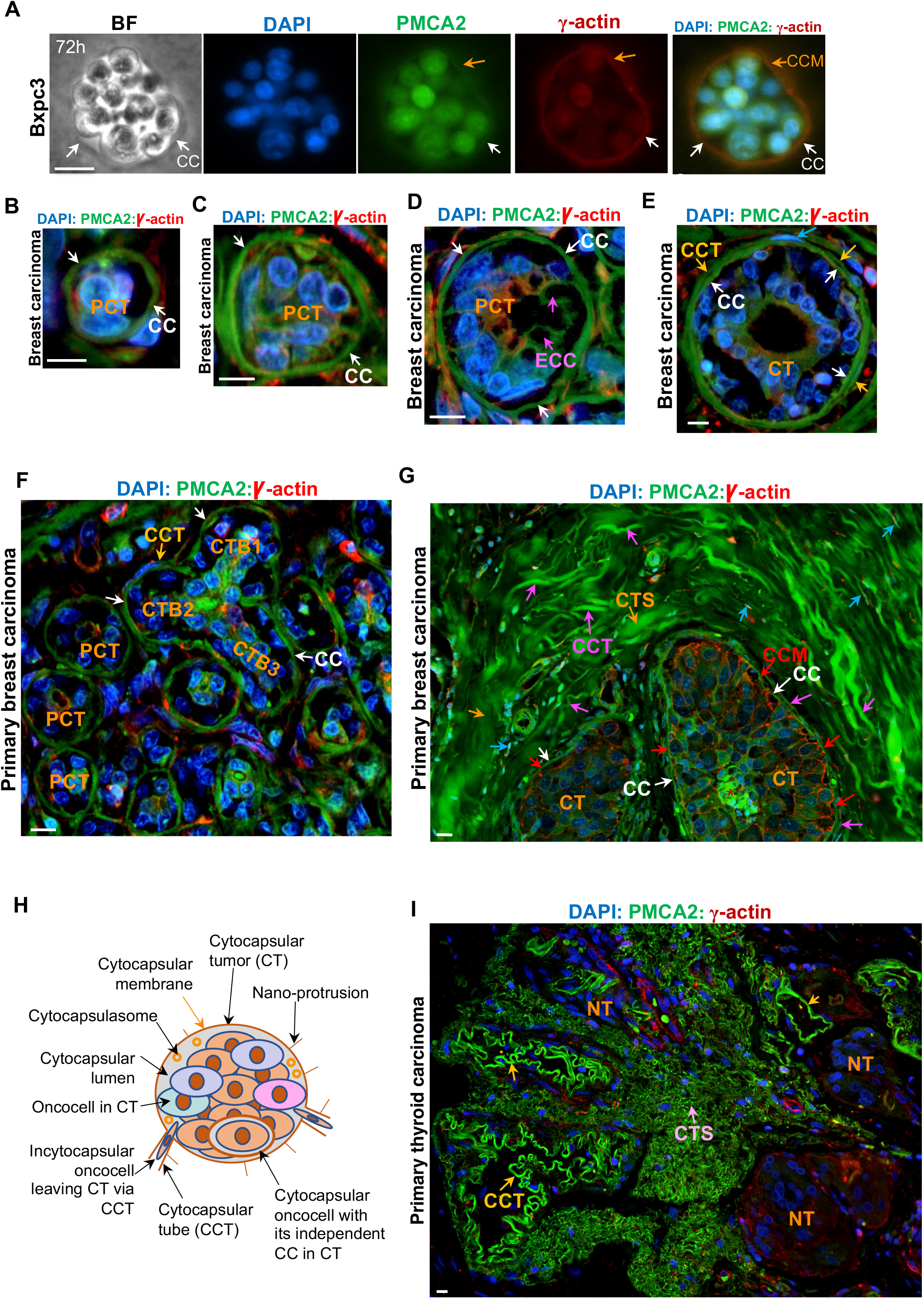
The lifecycle of cytocapsular tumor in human tissues. (**A**) Representative bright field (BF) and fluorescence microscope images of cytocapsular tumorspheres of Bxpc3 pancreas cancer cells in CC/CCT culture kit. Cytocapsula (white arrow) and cytocapsular membrane (orange arrows) are shown. (**B**) Representative IHC fluorescence microscope image of initiation-stage prophase cytocapsular tumor (PCT, <20 μm in diameter/width) in early breast carcinoma. (**C-D**) Representative IHC fluorescence microscope images of PCTs in early-development (C, <30 μm in diameter/width) and middle-development stage (D, (<40 μm in diameter/width) stages. Cytocapsula (CC, white arrows), ecellulated CC (ECC, purple arrows) are shown. ((**E**) Early stage cytocapsular tumor (CT) with cytocapsular tube (CCT, orange arrow) extended from CTs. (**F**) Representative IHC fluorescence microscope image of breast carcinoma tissues with high PCT and CT density. Multiple CTs merge into a bigger CT. A CT with 3 CT branches (CTBs), cytocapsula (CC, white arrow) and cytocapsular tube (CCT, orange arrow) are shown. (**G**) Representative IHC fluorescence microscope image of developed cytocapsular tumors (CTs) with thick CCT layers wrapping the CTs outside. Cytocapsula (CC, white arrow) of CTs, cytocapsular membrane (CCM, red arrow), cytocapsular tube (CCT, purple arrows), incytocapsular oncocells (cyan arrows), and CCT strand (CTS, orange arrows) are shown. (**H**) A schematic diagram of a cytocapsular tumor with its CCTs, nano-protrusion, and incytocapsular oncocells with/without their independent CCs. (**I**) Representative image of an acytocapsular oncocell mass-CC/CCT complex (AMCC). Nuperphase tumor (NT), CCT (orange arrow) and degraded CCT strands (CTS, pink arrow) are shown. Scale bar, 10μm.

Next, we investigated how cytocapsular oncocells progress into malignant tumors in human tissues *in vivo*. Initially, in the early clinical breast cancer tissues, single cytocapsular oncocells’ extracellular cytocapsulas grow into >15μm in diameter/width. Single breast incytocapsular oncocells proliferate and form small oncocell masses composed of several oncocells (n≥2 oncocells) in the CC lumens (n=675, **Fig.2B**). There are no CCTs derived from these small oncocell masses at this stage (n=675, **Fig.2B**). We term this kind of early phase of small oncocell mass enclosed in enlarged cytocapsula without CCTs as “prophase cytocapsular tumor (PCT)” (**Fig.2B**). Subsequently, CCs of PCTs grow up and increase in diameter/width, and breast incytocapsular oncocells continue to proliferate and form bigger compact breast oncocell masses enclosed in the enlarged CCs (n=312, **Fig.2C**). Sometimes, acellular cytocapsulas (n=161) appear in breast PCT lumens (**Fig.2D**), indicating that PCT incytocapsular oncocells generate secondary independent cytocapsulas, and these secondary cytocapsular oncocells can perform ecellulation and create ecellulated cytocapsulas (or groups). Subsequently, these ecellulated cytocapsulas in PCTs autodegradate into thin strands followed by decomposition and disappearance (n=78, **Fig.2D**). Breast oncocells in the enlarged CCs can extend parts of CC membranes, increase CC membranes in area, deform CC membranes, and form tube-shaped CCTs providing membrane-sheltered freeways for incytocapsular oncocell dissemination outside of CCs of the compact tumors. These CCT membranes are the extensions of CC membranes of PCTs *in vitro* (n=65, **fig. S3C**) and *in vivo* (n=117, **Fig.2E**). We termed this previously unrecognized single malignant tumor, which is enclosed in an enlarged, extracellular and membranous cytocapsula, and performs oncocell activities in the cytocapsular lumen, and generates extended cytocapsular tubes outside for oncocell dissemination, a “cytocapsular tumor” (**Fig.2E**).

Subsequently, uncontrolled breast incytocapsular oncocell proliferation produces more oncocells, forms tube-like structures in cytocapsular tumor (CT) lumens, and grows into bigger malignant tumors in the enlarged CC lumens (n=117, **Fig.2E**). Frequently, in the early stage, two or more adjacent small breast PCTs, CTs, or PCTs and CTs merge into larger PCTs (or CTs) via CC membrane contact, integration, degradation, and open connection formation, and form long or irregular-shaped cytocapsular tumorspheres enclosed in longer and bigger CCs (n=145, **Fig.2F**). Some newly merged CTs have three or more tumor branches (n=23, **Fig.2F**). Later, with uncontrolled oncocell proliferation, the merged CTs with tumor branches remodel and transit into spherical, oval or irregular compact CTs (n=132, **Fig. 2G**). Subsequently, breast CTs generate large quantities of CCTs outside for oncocell metastasis. Many long CCTs surround cytocapsular tumors (CTs) and form thick CCT layers enveloping CTs, and the measured thicknesses of CCT layers of CTs are 22∼510μm (n=132 CTs, **Fig. 2G**). These observations suggest that cytocapsular tumors with incytocapsular oncocells and CCTs provide two inherent physical and structural drivers for the two features of malignant tumors (cancers) in clinical observations: uncontrolled proliferation and metastasis (**Fig. 2, G and H**).

Subsequently, in the late cancer stage, the CC membranes of CTs degrade, decompose and disappear, leaving compact acytocapsular oncocell masses without CCs (n=814, **Fig. 2I**). We termed this kind of tumor that is composed of acytocapsular oncocell masses after CC degradation without enlarged CCs wrapping the tumor as “nuperphase tumor” (“nuper”, means “late” in *Latin*). Some acytocapsular oncocells in nuperphase tumors (NTs) regenerate many new, long, and highly curved CCTs (n=1021NTs). NT oncocells can invade into these CCTs via alloentry and disseminate, leaving decreased oncocell density in place (n=25, **Fig. 2I**). NT acytocapsular oncocell masses and the newly generated CCTs form a big acytocapsular oncocell mass-CCT complex (AMCC) (n=825, **Fig.2I**). The lifecycle of cytocapsular tumors includes 4 successive procedures: 1) single cytocapsular oncocells generate prophase cytocapsular tumors (PCTs) without CCTs, 2) generation of cytocapsular tumors (CTs) with many CCTs for cancer metastasis, 3) CC degradation engenders nuperphase tumors (NTs) without CC enclosing oncocell masses, 4) some acytocapsular oncocells in NT regenerate new CCs/CCTs or new small CTs and form AMCCs (**fig. S3D**).

### Distribution of cytocapsular oncocells and tumors in human tissues and organs

Cancers universally occur in most human organs and tissues (*1*). Next, we investigated the distribution of cytocapsular oncocells and tumors in human organs and tissues. In the examined 6 kinds of normal human organ tissues from breast, colon, liver, lung, prostate and stomach, we don’t find cytocapsular oncocells/tumors (**Fig. 3A**, n=14 patients). In the tested 38 subtypes of benign tumors (**Fig. 3A and table S1**, n=126 patients), most benign tumors do not present cytocapsular oncocells/tumors, while 13.4% of benign tumors display cytocapsular oncocells, indicating the transformation and cytocapsular oncocell occurrence in these benign tumors. In the tested 290 types/subtypes of cancers of 34 kinds of human organs/tissues (except hematologic cancers in blood, patient number, n=9,770; specimen number, n=9,958), 100% of tested cancers show cytocapsular oncocells (**Fig. 3**, **Table 1, and table S2 to 4**). These observations suggest that human normal and benign tumor tissues do not generate cytocapsular oncocells, and that human cancers universally engender cytocapsular oncocells. Hematologic cytocapsular oncocells appear in bone marrow, lymph node, spleen and thymus (**Fig.3B**, **Tables 1 and table S2 to 4**). There are many acytocapsular oncocells localized beyond CCs and CCTs, suggesting that acytocapsular oncocells coexist with cytocapsular oncocells *in vivo*. Cytocapsular oncocells exhibit 3 prominent features: 1) high abundance of PMCA2 in CC/CCT membranes, 2) CCTs are 3∼10μm in diameter/width and up to >3000μm in length in the sectioned specimens, and 3) localize in CCs or migrate in CCTs (**Fig.3, Tables 1, and table S2 to 4**). Cytocapsular oncocells’ CCTs in 283 subtypes of tested solid cancers display vast diversities in CCT density, morphologies, super-structures, degradation, interconnections, incytocapsular oncocell migration in CCTs, bunches, mixture with acytocapsular oncocells, reflecting the considerable heterogeneity of cytocapsular oncocells in solid cancers (**Fig. 3B**, **Table 1, table S2-4**). Cytocapsular tumors universally present in human solid cancers, but not in hematologic cancers in the blood (**Table 1, table S2 to 4**). The above observations suggest that cytocapsular oncocells are universally distributed in solid cancers and in hematologic cancers in bone marrow, lymph node, spleen and thymus, but not in normal or benign tumor tissues.

**Fig. 3.**
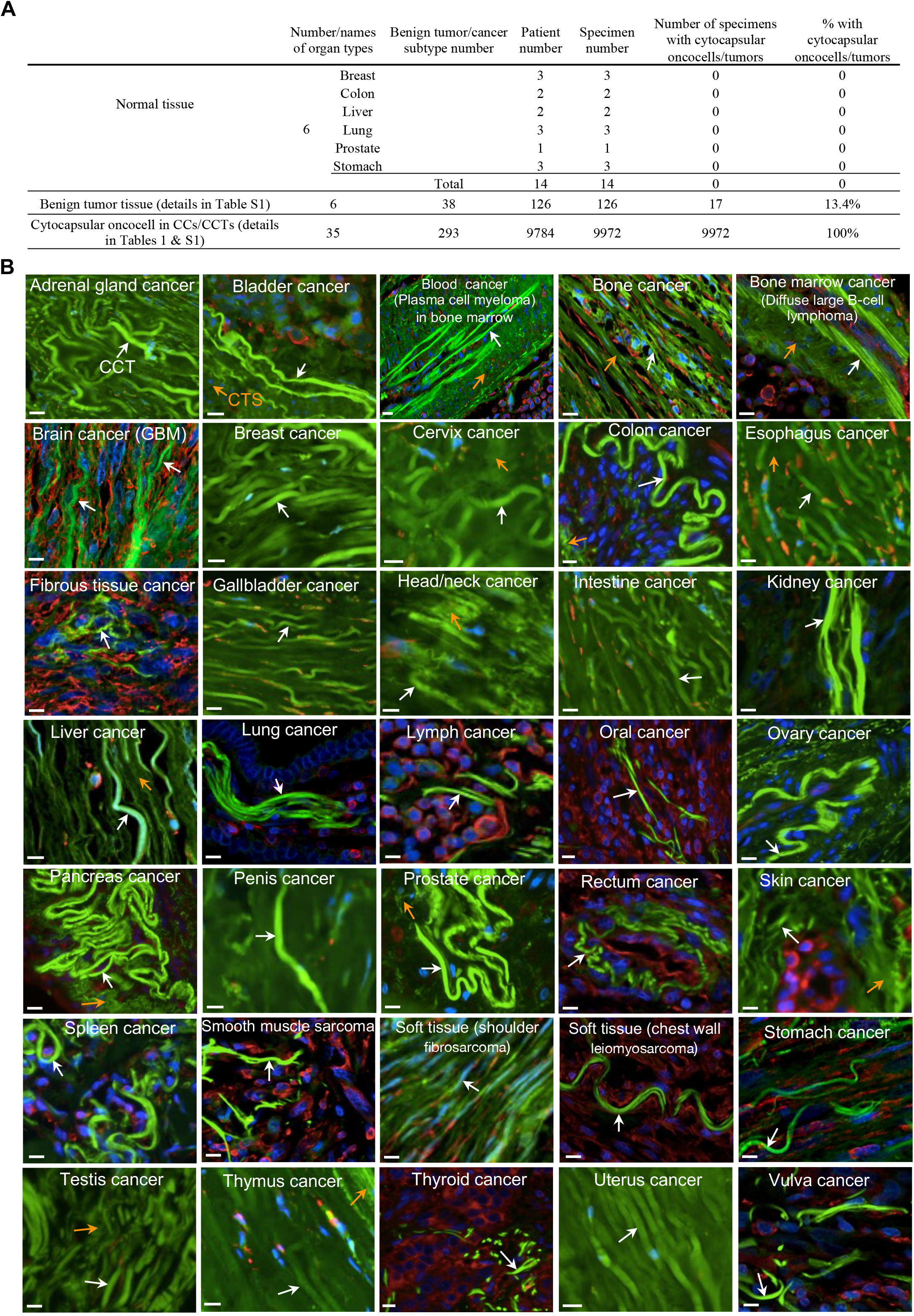
Distributions of cytocapsular tube tumors in 35 kinds of human tissues. (**A**) Table of statistical CCT presence in the checked 35 kinds of human tissues and organs. (**B**) Representative IHC fluorescence microscope images of CCTs (white arrows) in 35 kinds of human tissues and organs. Degraded CCT strands (CTS, orange arrows) are shown. Scale bar, 10μm.

**Table 1.**
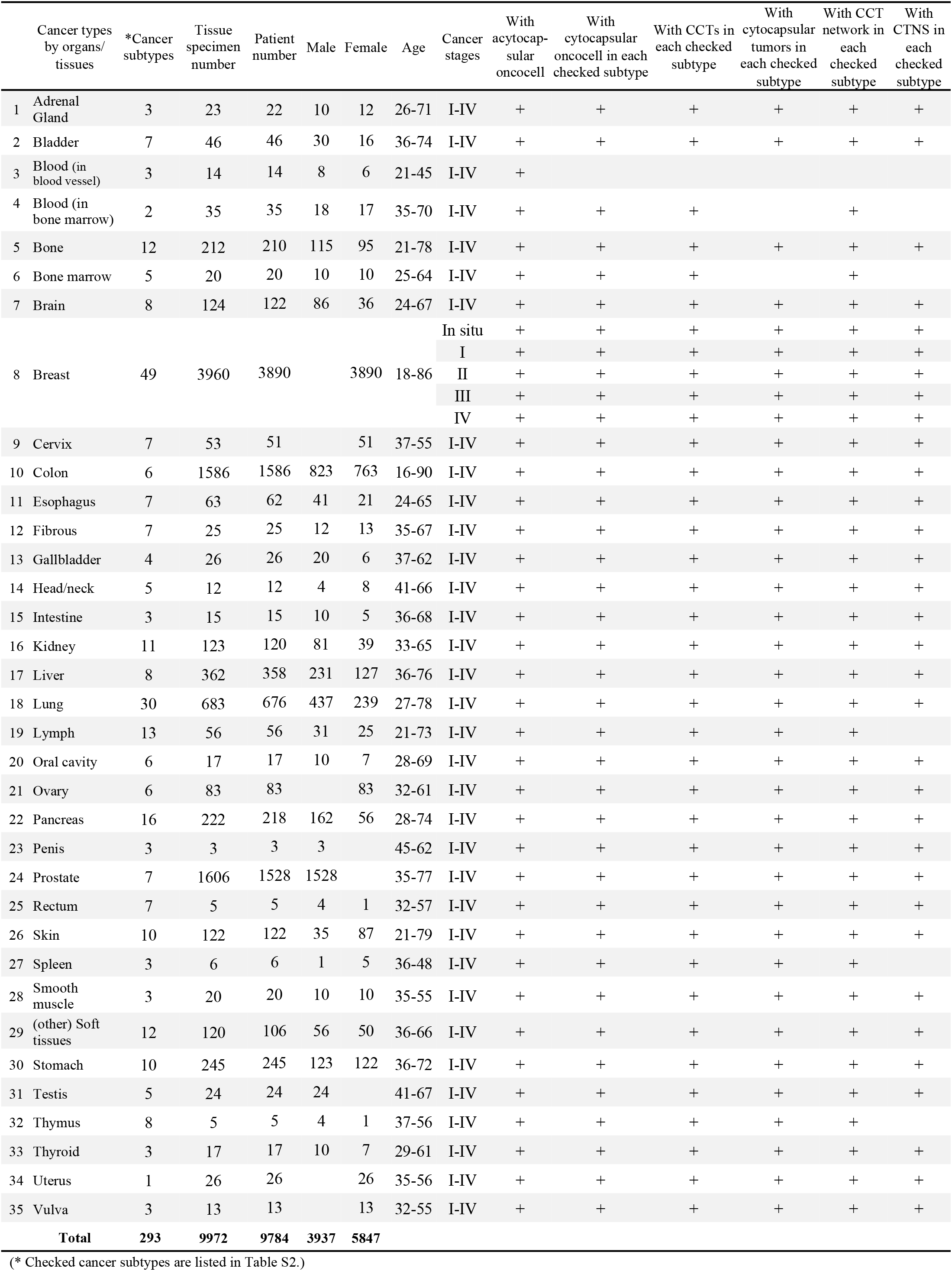
Characterization of oncocell, acytocapsular oncocell, cytocapsular oncocell, cytocapsular tumor, and cytocapsular tumor network system in 293 types/subtypes of cancers in human organs and tissues.

### Mechanics of cytocapsula growth, cytocapsular tube generation and elongation, and cytocapsular oncocell migration in CCTs

Next, we investigated the mechanics underlying cytocapsula growth of cytocapsular oncocells. Using inverted bright field microscope, we took videos of Bxpc3 tumorspheres in big CCs with wide circularity CC lumen gaps between tumorsphere edge and CC membranes. There are multiple spike-like structures (0.2∼0.9 μm in diameter/width, 4∼9μm in length) in the CC lumen, named as cytocapsular spike (CS, **fig.S4 and movie S1**). Cytocapsular spikes (checked CCs, n=26; average: n=4∼10 CSs per cytocapsular tumor) interconnect the tumorsphere surfaces and the CC membranes in multiple 3D directions, support the enlarged CC membranes, and maintain CC spherical/irregular morphologies without collapse (**fig. S4 and movie S1**). There are large quantities of tiny (0.1∼0.5μm in diameter/width), spherical, membrane-enclosed vesicles that attach to and cover the tumorsphere surfaces, indicating that tumorsphere oncocells generate large quantities of this kind of vesicles (n=12 cytocapsular tumorspheres, **fig. S4**). In the wide CC lumen gaps, there are some tiny free vesicles in constantly free and random motility formats in all directions in CC lumen fluids (**fig. S4 and 5, movie S1 and 2**). Many tiny vesicles detach from oncocell surfaces and freely move in the cytocapsular lumen fluids. Large quantities of tiny vesicles contact, fuse, and integrate into CC membranes, and increase CC membrane areas in size, and promote CC growth in size (**fig. S4 and 5, movie S1 and 2**). Consistently, there are many tiny, membrane-enclosed vesicles with high PMCA2 abundance located inside and outside of the cancer cell cytoplasm in both early-stage stomach cancer tissues *in vivo* (**fig. S6A**). Subsequently, these vesicles fuse together and generate cytocapsulas enclosing oncocells inside and engender cytocapsular oncocells (**fig. S6A**). These tiny vesicles supporting CC generation/growth are presented *in vitro* and *in vivo* (**fig. S4 and 5, and S6A**). We named this kind of tiny and membrane-enclosed vesicles with high PMCA2 abundance “cytocapsulasomes”, as they act as cytocapsular membrane building material delivery cargos and support CC generation and growth (**fig. S6**).

Cytocapsulasomes are initially engendered in the cytoplasm of acytocapsular transformed cells (n=56 samples, **fig. S6A**). Subsequently, cytocapsulasomes are released outside the cytoplasm membranes (n=87). These outside cytocapsulasomes fuse together and form small cytocapsular membrane fragments, which subsequently grow into cytocapsulas wrapping the whole oncocells and generate cytocapsular oncocells (n=62 cancer samples, **fig. S6A and movies S1 and 2**). The average detectable cytocapsulasome (CS) per cytocapsular oncocell in stomach and breast cancers are 34 ± 8 CS/cell (n=65) and 28 ± 7 CS/cell (n=120) *in vivo*, respectively (**fig.S6B**). The average detectable cytocapsulasomes in Bxpc3 cytocapsular tumorspheres with wide cytocapsular lumens is 126 ± 34 CS/CT (n=6) *in vitro* (**fig. S5B**). The above observations suggest that acytocapsular transformed cells initially generate cytocapsulasomes in the cytoplasm. Subsequently, cytocapsulasomes are released outside of cellular membrane and attach the cell membrane. Then, cytocapsulasomes gradually merge together to form cytocapsulas, and engender cytocapsular oncocells. Cytocapsulasomes function as cytocapsular membrane material delivery cargos to drive cytocapsula growth (**fig. S4 to 6**).

Next, we assessed the mechanics underlying cytocapsular oncocell CCT generation and elongation. Using time-lapse DIC microscopy, we investigated cytocapsular oncocell CCT generation and elongation *in vitro*. Initially, single MCF-7 breast oncocells generate CCs which tightly wrap themselves. With bleb-based motilities, single cytocapsula oncocells in CCs push CC membranes forward, deform CC membrane shapes into tube-like morphologies, and generate short CCT fragments (**fig.S7A, 1 to 3, and movie S3**). The initial CCT fragments are well anchored in the viscous CC/CCT culture kit matrix and maintain the wide tube shapes without being stretched and contracted into thin and long CCT tail shapes (**fig.S7A, 2 and 3, and movie S3**). The cytocapsular oncocells constantly generate cytocapsulasomes, which merge and integrate into the front part of the CC membranes that tightly contact with oncocell membranes, and increase CC membrane areas. Cytocapsular oncocells constantly and dynamically generate many transient blebs in various sizes. Blebs repeatedly protrude and retract in the CC, sense the extracytocapsular environments in many 3D directions, and choose and decide the motility directions. After cytocapsula creation, oncocells migrate backward in CCTs they generated. They engender several short cytocapsular spikes in the CCT lumens, which link the rear of the oncocell and CCT membranes (**fig. S7A**). Subsequently, when the single cytocapsular oncocells migrated backward in the established CCTs, it switched into a lamellipodia-based motility format (**fig. S7A, 11 to 20 and movie S3**). Average CCT elongation speed of MCF-7 cytocapsular breast oncocells in CC/CCT culture kit matrix is 1.25 ±0.3μm/min (n=3 cells, **fig. S7B**). Next, we investigated cytocapsular oncocell CCT generation and elongation *in vivo*. In the early stage of breast cancer, there are many initial cytocapsular oncocell CCTs that have long and thin CCT tails (0.1∼1μm in width), indicating that the initial CCT fragments are stretched and contracted by the single cytocapsular breast oncocells when they move forward and the initial CCT fragments are not well anchored into the ECM (**fig. S7C**). Subsequently, when the CCTs are well anchored by CCT nano-protrusions extended into the ECM, the CCTs are consistently maintained at 3∼6μm in diameter/width in solid cancers (**fig. S7C**). The above *in vitro* and *in vivo* observations suggest that single cytocapsular oncocells in CCs employ a bled-based sensing and motility format to choose directions and move forward, engender cytocapsulasomes to support CC membrane size increase, push and deform CC membranes into tube-shaped morphologies, and generate and elongate CCTs (**fig. S7D**). The initial CCT fragments frequently have long and thin CCT tails *in vivo*.

Next, we investigated mechanics of cytocapsular oncocell migration in CCTs. Single Bxpc3 pancreas cytocapsular oncocells in migration in long CCTs (3∼6μm in diameter) are suppressed by the stretched and contracted CCT membranes and shaped into long, thin, spindle-shaped morphologies. Single cytocapsular oncocells are usually in single cell mesenchymal migration in CCTs (**fig. S7E**). Furthermore, using time-lapse DIC technologies, we investigated the dynamic cell migration activities in CCTs with multiple MCF-7 breast cytocapsular oncocells in migration in a long CCT. In most instances, multiple cytocapsular oncocells in CCTs are in a single epithelial migration format, not in collective migration mode. The polarized, thin and long cytocapsular oncocells in CCTs migrate forward with periodic protrusion and retraction of the leading lamellipodia and movement of the cell rear. Sometimes, the long lamellipodia are completely retracted and the cells transiently appear in spherical morphologies in CCTs. CCT membranes always tightly wrap and adhere to cancer cell cytoplasm membranes and dynamically increase/decrease the CCT diameter/width locally, displaying considerable CCT membrane elasticity (**fig. S7F and movie S4**). The polarized cytocapsular oncocells in CCTs can freely switch the migration direction into an opposite direction and therefore can migrate bi-directionally in CCTs. Here, multiple cytocapsular oncocells freely and bi-directionally migrate in a long CCT, with dynamic cellular morphologies. The elastic CCT membranes shelter from obstacles in the heterogeneous extracytocapsular matrix outside, and provide membrane-enclosed and protected freeways for cytocapsular oncocell migration inside (**fig. S7, E and F, and movie S4**). The average cell migration speed of multiple single Bxpc3 pancreas cytocapsular oncocells in CCT *in vitro* is 2.7 ± 0.5μm/min (n=10 cells (**fig. S7G**). Consistently, in the long (straight or curved) cytocapsular colon oncocell CCTs (3∼6μm in diameter) in colon carcinoma tissues, cytocapsular colon cancer cells in migration in CCTs are usually in single, long, thin and spindle-shaped morphologies *in vivo* (CCT, n=256; tissues, n=34; **fig. S7H**). This indicates that cytocapsular oncocells in migration in CCTs *in vivo* use a single mesenchymal migration format (**fig. S7H**). The above observations suggest that cytocapsular oncocells can freely, dynamically, and bi-directionally migrate in membrane-enclosed and protected CCTs in a single mesenchymal migration format and in thin and long morphologies *in vitro* and *in vivo*. A CCT with multiple or numerous oncocells inside is topologically and bio-functionally a long and tube-shaped cytocapsular tumor, and is thus termed a “cytocapsular tube tumor” (**fig. S7, H and I**).

### Primary cytocapsular tumor network system progression

Primary malignant tumors are origins of metastatic (secondary) cancers. Thus, we asked how primary cytocapsular tumors progress in primary niches. Then we assessed cytocapsular tumorsphere metastasis and secondary cytocapsular tumorsphere growth *in vitro*. In the CC/CCT culture kit matrix, at 68h, single Bxpc3 cytocapsular oncocells proliferate in CCs and grow into big primary cytocapsular tumorspheres (CTs, **fig. S8A**). Two or more primary CTs merge together via CC membrane contact, degradation, and open connection formation, and form long and irregular-shaped CTs enclosed in long and big CCs (**fig. S8A**). Cytocapsular oncocells in enlarged CCs push CC membranes, elongate and generate CCTs. CCTs interconnect and form CCT networks (**fig.S8A**). The CCT networks interconnect all primary and secondary CTs and form a cytocapsular tumorsphere network systems (CTNS) in a well of 6-well plate (**fig. S8A**). Acytocapsular oncocells can enter CCTs via alloentry. Incytocapsular oncocells perform spontaneous ecellulation and become acytocapsular oncocells beyond CCs/CCTs (**fig. S8A**). The ecellulated acytocapsular oncocells can regenerate new cytocapsulas and CCTs (**fig. S8B**). Incytocapsular oncocells metastasize via CCT networks (**fig. S8B**). In addition, cytocapsular oncocells metastasize and reside in CCT interconnection nodes, proliferate and grow into secondary cytocapsular tumorspheres, which are integrated into the established CTNSs (n=38 checked secondary cytocapsular tumorspheres, **fig. S8, A and B**). Incytocapsular oncocells migrate and translocate in CTNSs via CCT networks (**fig. S8B**). Cytocapsular tumorspheres’ spontaneous ecellulation engenders acellular CT cytocapsular parts and acellular CCT fragments, making the open CCT connections between CTs visible (**fig. S8B**). At 78h, all the primary and secondary CTs in a well of 6-well plates are interconnected and covered by the integrated cytocapsular membrane systems, which are composed of cytocapsular tumorsphere CCs and CCT networks (CTNS number, n=55, **fig. S8C**). These observations suggested that: (1) primary CT metastasize, generate CCTs, CCT networks and primary CTNSs; (2) metastasis of cytocapsular oncocells develop into secondary cytocapsular tumorspheres, and form secondary CTNSs; (3) ecellulation and alloentry allow oncocells to bidirectionally evict from and enter into CCs/CCTs; and (4) primary and secondary CTNSs integrate via CCT networks and form a dynamic and integrated primary and secondary CTNS enclosed in cytocapsular membrane systems *in vitro*.

Next, we investigated primary cytocapsular tumor network systems (CTNSs) *in vivo*. *Prophase CT (PCT) formation*: In early-stage primary breast cancers, there are many spherical or irregular-shaped prophase cytocapsular tumors (PCTs) in variable sizes (25∼120μm in diameter/width, **Fig. 2F and fig. S9A**). The average prophase cytocapsular tumor (PCT) densities of early-stage cancers in breast, colon, and prostate are 202 ± 59 PCTs/mm^2^, 125 ± 32PCTs/mm^2^, and 173 ± 26PCTs/mm^2^, respectively (specimens, n=3∼6, 1 specimen/patient, 1-2 subtypes/cancer type, **fig. S9B)**.

*PCT to CT:* Subsequently, the PCTs develop into spherical or irregular-shaped cytocapsular tumors (CTs) in variable sizes (50∼320μm in diameter/width, **Fig. 4A and fig. S9C**). The average CT densities of early-stage cancers in breast, colon, and prostate are 176 ± 38CTs/mm^2^, 87 ± 34CTs/mm^2^, and 158 ± 28CTs/mm^2^, respectively (specimens, n=5∼12, 1 specimen/patient, 3-4 subtypes/cancer type) (**fig.S9D**). These data indicate that PCT and CT densities in the early primary solid cancers are statistically high. Consistent with cytocapsular tumorsphere merge *in vitro*, two or more small PCTs/CTs *in vivo* can merge into middle or big sized PCTs/CTs (**Fig. 2F and 4A**). The CTs with >50μm in diameter/width start to generate a few cytocapsular tubes (CCTs) outside. Cytocapsula tumors with >70 μm in diameter/width engender a lot of CCTs outside and form thick surrounding CCT layers wrapping cytocapsula tumors (**Fig. 2G and 4B)**. The dense CCT layers present various thicknesses (30∼803μm in thickness) (**Fig. 4B**). Straight, curved or coiled CCTs intensively interconnect and form 3D CCT networks, which broadly interconnect primary CTs in the primary cancer niches (**Figs. 4B and fig. S9E**). These observations suggest that cytocapsular tumors in the primary niches are physically interconnected via 3D CCT networks and form primary cytocapsular tumor network systems (CTNSs) *in vivo*.

**Fig. 4.**
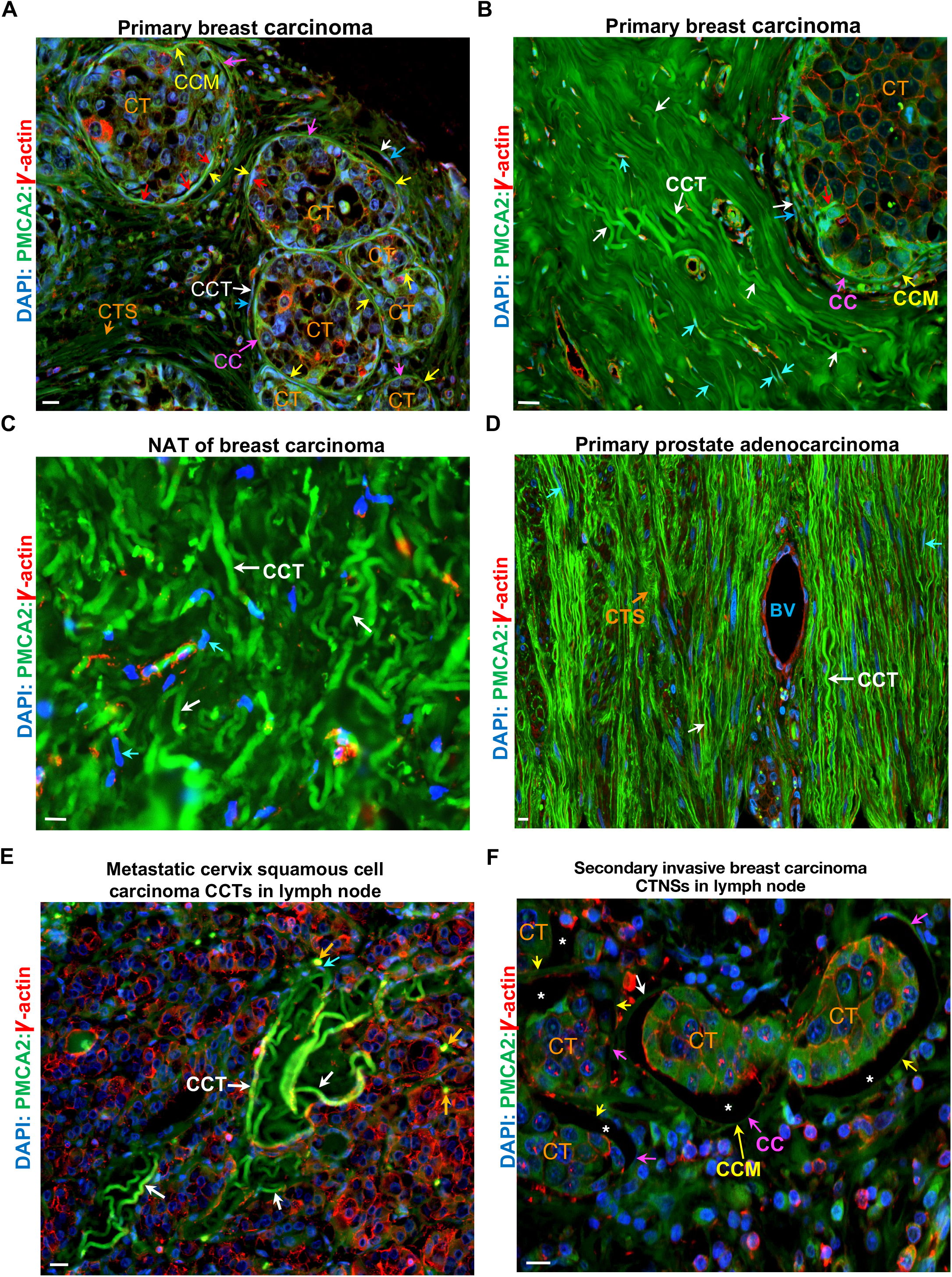
Primary and secondary cytocapsular tumor network system progression. (**A**) Enlarged image from Fig.S9C. Multiple cytocapsular tumors (CTs) interconnect by CCT networks and form cytocapsular tumor network systems (CTNSs) in primary breast carcinoma. Cytocapsula (CC, purple arrows) and cytocapsular membrane (CCM, yellow arrows) of CTs, cytocapsular tube (CCT) outside CT (white arrows), and inside CT (red arrows), incytocapsular oncocells in migration in CCTs (cyan arrows), and degraded CCT strand (CTS, orange arrow) are shown. (**B**) Representative image of a cytocapsular tumor (CT) with a thick CCT layer wrapping the CT. Cytocapsula (CC, purple arrow) and cytocapsular membrane (CCM, yellow arrow) of CT, cytocapsular tube (CCT, red arrow) generated by oncocells in CTs, CCTs outside the CT (white arrow), incytocapsular oncocells (cyan arrows) in migration in CCTs outside of CTs are shown. (**C**) Representative image of CCT network masses in “normal tissues adjacent to tumor(NAT)”. CCTs (white arrows) and incytocapsular oncocells in migration in CCTs (cyan arrows) are shown. (**D**) Representative image of primary malignant prostate adenocarcinoma tissues with cancer metastasis, which harbor blood vessel (BV) and large quantities of CCTs with oncocells in migration in CCTs. CCTs (white arrows), CCT strand (CTS, orange arrow), and incytocapsular oncocells in migration in CCTs (cyan arrows) are shown. (**E**) Representative image of metastatic cervix squamous cell carcinoma CCTs (white arrow) invaded into lymph node and coexist with massive number of immune cells. Exposed surfaces of perpendicular cross-sectioned CCTs (orange arrows) are shown. (**F**) Representative image of secondary invasive breast carcinoma cytocapsular tumor network systems (CTNSs) in lymph node. Cytocapsular tumor (CT), cytocapsula (CC, purple arrow) and CC membrane (CCM, yellow arrows), and wide C-shape or irregular shape cytocapsular lumens in the sectioned CTs (white asterisk), and CCTs (white arrow) are shown. Scale bar, 10μm.

*CC degradation*: Subsequently, CCs and CCTs of cytocapsular tumors in size of >800μm in diameter/width degrade into strands followed by disappearance, engendering acytocapsular malignant tumors/oncocell masses with high cell density and without CCs or CCTs (**fig.S9F**). *Nuperphase Tumor*: Uncontrolled proliferation of acytocapsular oncocells generate big and irregular-shaped acytocapsular malignant tumors/oncocell masses (up to >2cm in width) in the checked sectioned specimens (**fig. S9G**).

*AMCC*: The acytocapsular status of these big malignant tumors/oncocell masses is transient. Subsequently, some acytocapsular oncocells regenerate a few or many new straight, curved or coiled CCTs, and form CCT network bunches and masses, which invade into and scatter in the dense acytocapsular oncocell masses, making a complex mixed by dense oncocell masses and CCT networks but without an enlarged CC wrapping them (**fig.S9H**). CCT bunches and masses create many CCT mass cavities (cross section: in round, oval or irregular shapes, 30∼200μm in diameter/width; longitudinal section: in straight, curved, or irregular column morphologies, 80∼850μm in length), which are interconnected and form CCT mass cavity compound (**fig. S1C3 and S9H**). Acytocapsular oncocells around CCTs perform alloentry, invade into CCTs, and disseminate via CCTs (**fig. S9H**).

*AMCC with CCT degradation*: Subsequently, CCTs degrade and disappear, leaving CCT mass cavities filled with intercellular fluids (**fig. S9I**). Meanwhile, some acytocapsular oncocells regenerate new CCTs (**fig. S9J**). CCT degradation and regeneration coexist in the Acytocapsular oncocell Mass-CCT network Complex (AMCC) (**fig. S9J**).

*Post-AMCC*: Subsequently, after many oncocells enter CCTs and depart away via CCTs, there are only a few oncocells left, and many CCTs (**fig. S9J**). Then, CCT degrade, decompose and disappear (**fig. S9K**). CTNSs in primary niches at the late cancer stage still remain (**fig. S9L)**. Some small or middle sized CTs present oncocell apoptosis and CCT degradation inside (**fig. S9L**). The above observations suggest that: 1) primary CTs develop into primary CTNSs *in vivo*, 2) CC/CCT degradation, acytocapsular oncocell proliferation, and CC/CCT regeneration generate AMCC, 3) primary CTNSs are dynamic systems including CCT alloentry, ecellulation, CC/CCT generation, degradation and regeneration, and dissemination of cytocapsular oncocells in CCTs.

Normal tissues adjacent to the tumor (NAT) are necessary sites that CCTs have to go through if they expand beyond. Indeed, there are large quantities CCTs that aggressively invade into and go through the NATs in one or multiple 3D directions in breast cancer tissues (**Fig. 4C and fig. S10A**), many CCTs in the bone marrow NAT of plasma cell myeloma (**fig.S10B**). Even in the hard tissue NAT of trabecular bone of plasma cell myeloma, there are a few CCTs (**fig.S10C**). In the examined NATs of 68 subtypes of cancers, 100% of them harbor a lot of CCTs and networks with cytocapsular oncocells in migration inside (**fig. S10D**). CCT density in soft tissue cancers can be up to 114CCT/mm^2^ and even in hard tissue (bone) cancers can be up to 10CCT/mm^2^ (**fig. S10D**). The above observations (**Fig 4, B and C, fig. S9 and 10, Tables 1, and table S1 to 4**) suggest that cytocapsular tumor progression in the primary niches includes 7 major successive stages: 1) generation of transformed acytocapsular oncocells, 2) acytocapsular oncocell devolution and generation of cytocapsular oncocells, 3) generation of prophase cytocapsular tumors and cytocapsular tumors, 4) formation of CCT networks and primary CTNSs, 5) cytocapsular oncocell metastasis via CCTs and go through NATs, 6) CC/CCT degradation and AMCC, 7) CCT regeneration, alloentry, ecellulation, and dynamic CTNS formation with continuous cytocapsular tumor metastasis ad CTNS regeneration.

### Cytocapsular oncocell metastasis in human tissues *in vivo*

Cancer metastasis is a major source of cancer lethality (*15–17*). Next, we assessed how cytocapsular oncocells in CCTs disseminate across diverse tissues and organs *in vivo*. Single CCTs invade into and spread not only in loose (**fig. S11A**) and compact (**fig. S11, B and C**) tissues, but also in hard tissues of trabecular bone (**fig.S10, C and D**), suggesting that cytocapsular oncocell in single CCTs can invade into and go through various kinds of tissues with diverse cell densities and ECM/matrix hardness. Single CCTs can be highly curved and very long (**fig.S11D**). 3D CCT networks enhance collective cytocapsular oncocell metastasis in and cross various kinds of tissues (**fig.S9, E and F, fig. S11, E and F**). Massive and compact CCT network bunches increase metastasis freeway density and elevate cytocapsular oncocells dissemination efficiency (**fig. S11, G to K**). Super-large structures of CCT networks facilitate the acytocapsular oncocells in the compact tumors/oncocell masses to enter CCTs and disseminate via CCTs (**fig. S11L**). Cytocapsular oncocells and CCT networks presented in bone marrow (**fig.S10B, Tables 1, table S2 and 3**), lymph nodes (**Tables 1, table S2 and 3**), spleen, and thymus (**Table 1, table S2 and 3**) suggest that CC/CCT membranes can effectively shelter immune cells and their attacks outside, and therefore protect cytocapsular oncocells inside. CCT networks are presented in 290 subtypes of cancers and in 34 kinds of human tissues (**Fig. 3 and 4, fig. S9 to 11, Tables 1, table S2 and 3**), suggesting that CCTs effectively cross all kinds of human tissues and organs for cytocapsular tumor metastasis. Large quantities of CCTs are found outside humoral vessels (**Fig. 4D, fig. S11, H to K, fig. S12A**), and that in the primary niches, NATs, and secondary niches, the average ratios of the density of CCTs to that of humoral vessels are up to 289 ± 6 folds, such as in **fig. S12B**. The above observations suggest that cytocapsular tumor metastasis via CCT freeway systems dominates tumor metastasis *in vivo*. Occasionally, a very low percentage of CCTs invade into micro blood vessels and release oncocells into the circulation systems (**fig. S12, C to F**), indicating that humoral vessel CCT invasion-caused oncocell release is a source of circulating tumor cells in the blood. Furthermore, we tested metastases of cytocapsular oncocells in 35 types of cancers (1∼10 secondary niches per type), and observed that cytocapsular tumors broadly disseminate to multiple secondary niches (**Table 1 and table S3**), which are consistent with clinical observations that primary tumors always metastasize into multiple tissues and organs.

### Secondary cytocapsular tumor network system progression

Cancer metastasis and secondary tumors caused multiple tissues/organs biological function failure are major sources of cancer lethality^8^. Thus, we assessed if metastasized cytocapsular oncocells in secondary niches generate secondary cytocapsular tumors and CTNSs *in vivo*. Indeed, after breast cytocapsular oncocell CCT networks metastasize and arrive at lymph nodes, they initially form thin CCT layers wrapping lymph nodes (**fig.S13A1**). Subsequently, with CCT branching morphogenesis, more breast cytocapsular oncocell CCTs are generated and form thick CCT layers, enveloping lymph nodes (**fig.S13A2**). Cervix cytocapsular oncocell CCTs massively invade into lymph nodes with compact lymphocytes, and a lot of metastasized-cervix cytocapsular oncocells disseminate into dense lymph nodes (**Fig. 4E and fig. S13B**). Subsequently, in lymph nodes, metastasized breast cytocapsular oncocells in CCTs proliferate, and generate many small secondary breast cytocapsular tumors in enlarged CCs (**Fig. 4F and fig. S13C**). In the early stage, secondary cytocapsular breast tumors have C-shaped, circularity or irregular-shaped CCT lumen gaps (up to 30μm in width) between cytocapsular oncocell mass surface and CC membranes (**Fig. 4F and fig. S13C**). Breast cytocapsular oncocells in CCTs metastasized at lymph nodes grow into large quantities of secondary breast cytocapsular tumors, which occupy the spaces of normal lymph node cells and many normal lymph node cells disappeared (**fig. S13C**). These secondary breast cytocapsular tumors are interconnected via CCT networks and form secondary breast CTNSs in the secondary niches (**fig. S13C**). The average densities of the checked secondary breast cytocapsular tumors in bladder, liver and lymph node are 163 ± 71CTs/mm^2^, 160 ± 68CTs/mm^2^, and 170 ± 56CTs/mm^2^, respectively (patients, n=4∼6) (**fig. S13D**). The above observations (**fig. S9 to 11 and S13, A to D**) indicate that the primary CTNSs are physically interconnected with secondary CTNSs via CCT networks and form integrated primary and secondary CTNSs.

Small nasopharynx secondary cytocapsular tumors grow up and engender many CCTs (**fig.S13E**). CCs and CCTs of ovary secondary cytocapsular tumors in omentum degrade, and generate big acytocapsular ovary tumors/oncocell masses (up to 2cm or more in diameter/width) without CCs/CCTs (**fig. S13F**). Subsequently, some oncocells in acytocapsular ovary tumors in omentum regenerate new CCTs CTs, and CNTSs (**fig. S13G**). Consistently, some cervix oncocells of acytocapsular cervix tumors in lymph node engender many CCTs in lymph nodes (**fig. S13H**), and rectum oncocells in acytocapsular rectum tumors in mesentery engender large quantities of new CCTs and networks in mesentery (**fig.S13, I and J**). Acytocapsular oncocells invade into the regenerated CCTs via alloentry and leave away (**fig. S13, G to J**). Many secondary colon acytocapsular oncocells in liver disseminate via regenerated colon CCTs, leaving many spaces without cells and only filled with intercellular fluids (**fig. S13K**).

In the late cancer stage, after secondary dissemination of metastasized cytocapsular tumors via CCT networks, CCTs degrade, and CCT networks decompose (**fig. S13, K and L**). Sometimes, many red blood cells randomly spread in large areas in the secondary hepatocellular carcinoma in cerebrum of brain, indicting some (micro)blood vessels are broken and leaky caused by CCT invasion and (micro)blood vessel decomposition, and red blood cells are released (**fig.S13L**). The above observations suggest that metastatic cytocapsular tumor progression in the secondary niches includes 6 major successive stages: 1) arrival and invasion of metastatic cytocapsular oncocells in CCTs in the secondary niches, 2) generation of secondary cytocapsular tumors, 3) formation of secondary CTNSs, 4) formation of dynamic integrated primary and secondary CTNSs via CCT networks, 5) CC/CCT degradation and formation of AMCC, 6) generation of new CCTs for oncocells’ next metastasis (**fig.S13, table S2 to 5**). These observations (**fig. S9 to 13, Tables 1 and table S2 to 5**) suggest that metastasized cytocapsular oncocells have capacities to generate large quantities of small/middle/big-sized secondary cytocapsular tumors and dense CTNSs in the secondary niches (in neighboring or far-distance organs/tissues), and lead to massive normal cell disappearance followed by affected, harmed, or even failed biological functions in the secondary niches related tissues/organ. In summary, our results suggest that cytocapsular oncocells, cytocapsular tumors, and integrated primary and secondary CTNSs coordinate membrane-sheltered cancer progression in human (**Fig. S14**).

## Discussion

The mechanisms of cancer progression in human are critical for cancer research, early screening, prognosis, diagnosis, drug development, therapy and treatment (*8–11*, *18–20*). Here we investigated mechanisms of cancer progression in human tissues and organs *in vivo*, and found that cytocapsular oncocells, cytocapsular tumors, and cytocapsular tumor network systems drive membrane-sheltered cancer development and progression.

The identification of PMCA2 as a cytocapsular membrane protein marker, which makes the previously invisible CCs/CCTs visible and facilitate to understand cytocapsular oncocells/tumors with newly discovered organelles/compartments, may open new avenues for cancer research and therapies. The findings of the cytocapsular oncocell, cytocapsular tumor, and cytocapsular tumor systems (CTNSs) unveil a long-term unrecognized mechanism that cytocapsular membrane systems function as integrated, membrane-encompassed, and protective powerhouses coordinating cytocapsular oncocell proliferation, cytocapsular tumor growth, cytocapsular cancer metastasis via and in CCT networks. Cytocapsular oncocell metastasis in cytocapsular membrane-enclosed integrated primary and secondary CTNSs has several advantages: 1) free of obstacles from heterogeneous extracytocapsular matrix and neighboring cells, 2) minimum or free of extracytocapsular attacks from immune cells and stresses from toxic molecules (such as cancer drugs), 3) have membrane-enclosed physical freeway systems for safe and efficient cancer metastasis, 4) can perform bi-directional migration in the dynamic and integrated primary and secondary CTNSs.

The finding of integrated primary and secondary CTNSs may shed light to cancer researches to treat with the primary and secondary tumors into a united, dynamic and cytocapsular membrane-encompassed system, not in a fragmental view of isolated or separated primary and secondary tumors that clinically display transient and limited success followed by irreversible and undruggable cancer relapse. The finding of the combined extracellular membranes of cytocapsular oncocells, cytocapsular tumors, and primary and secondary CTNSs may facilitate the researches for the development of better pharmaceutical and immune therapies with less or minimum membrane barrier caused pan-drug resistance and immune attack escape. The occasional CCT invasion into humoral vessels and oncocell release as a circulating tumor cell resource are consistent with clinical observations that even massive tumor metastases occur while the circulating tumor cells are in very low percentage in blood cells *in vivo*. Cytocapsular tubes can invade into and pass through loose and compact tissues, soft and hard tissues/ECM, and hard matrices (bone), which is consistent with the clinical observations that solid tumors can disseminate into most of human tissues and organs including bone metastases. The capacities of primary and multiple secondary devolution of some oncocells and generation of new cytocapsular tumors and CCTs are in agreement with clinical observations of multiple secondary cancer metastasis and multiple occurrences of tumor relapse during cancer therapies. Our study raises a question of what are the molecular mechanisms underlying CC/CCT initiation of transformed cells *in vivo*.

Our findings reveal cytocapsular oncocell, cytocapsular tumor, cytocapsular oncocell metastasis in CCT networks, and cytocapsular tumor network systems in cancer development and progression, which may facilitate the researches for effective therapies against cancers.

## Acknowledgments

We greatly acknowledge Dr. Ed Harlow of Harvard Medical School (USA) and Cancer Institute of University of Cambridge (UK), Dr. Nahum Sonenberg of McGill University of Canada, Dr. Michael Roehrl of Beth Israel Deaconess Medical Center (BIDMC), Dr. Stephen Ganshirt of Northwest University Medicine Hospital, Dr. Joan Brugge, David Golan and Mark Namchuk of Harvard Medical School, Dr. Yubo Yang and Dr. Qiping Hou for their help and meaningful discussion in the study. **Funding**: Supported by a grant (to Dr. Yi) from Cytocapsula Research Institute Fund for Cytocapsula Research, and a grant (to Dr. Yi) from Centiver Ltd Fund for Cytocapsula Research. All data produced in the present work are contained in the manuscript. **Author contributions**: T.Y. designed and performed experiments, collected and analyzed data, and wrote manuscript. G.W. took part in data analyses and manuscript writing. **Competing interests:** No competing interests. **Data and materials availability:** All data and materials are available.

## Supplementary Materials

Materials and Methods

Supplementary Text

Figs. S1 to S14

Tables S1 to S5

References (*22–23*)

Movie S1-S4

