## Supplementary material for "Malignant tumors engender and employ cancer-specific organelles for self-protection and metastasis": supple text

**Materials and Methods**

**Reagent, antibodies and devices**

CC/CCT culture kits (Cat. CD0104, 6-well plates; Cat. CD0105, 12-well plates, and Cat. CD 0106, 24-well plates) and kits with glass cover slips in the well bottom and embedded by CC/CCT culture matrix layer (Cat. CD 0112, 6-well plate) were ordered from Celldevi Inc. CC/CCT fixation kit (CD0201) were ordered from Celldevi Inc. Cancer cell lines of pancreas cancer cell Bxpc3, breast cancer cell MCF-7 and colon cancer cell SK-CO-1 and cell culture media were ordered from ATCC. ^13^C_6_, ^15^N_2_-L-Lysine (Cat. 88209) and ^13^C_6_, ^15^N_4_-L-Arginine (Cat. 89990) for SILAC labeling were ordered from Thermo Fisher Scientific. Unipick^TM^ and capillary units were ordered from NeuroInDx Inc. Rabbit anti-PMCA2 antibody (polyclonal, ab3529; 1:200 dilution), mouse anti-γ-actin antibody (monoclonal, ab123034; 1:200 dilution ) for immunofluorescence assay were ordered from Abcam. DAPI (1:1,000 dilution in the immunofluorescence assay) was ordered from KPL. Human normal and cancer tissue specimens were ordered from TissueArray and US Biolab (or gifted by local hospitals).

**Stable SILAC labeled CC/CCT culture and collection for CC/CCT proteome analyses**

Following CC/CCT culture kit usage instructions, stable SILAC labeled cancer cells of Bxpc3, MCF-7 and SK-CO-1 with ^13^C_6_, ^15^N_2_-L-Lysine and ^13^C_6_, ^15^N_4_-L-Arginine were implanted in CC/CCT culture kits (6-well plates) with cell culture media with ^13^C_6_, ^15^N_2_-L-Lysine and ^13^C_6_, ^15^N_4_-L-Arginine^1^. Stable SILAC labeled cancer cells generate stable SILAC labeled cytocapsulas (CCs) and cytocapsular tubes (CCTs). Sometimes, some incytocapsular oncocells are spontaneously evicted from CCs and CCTs *in vitro*. After ecellulation, evicted cancer cells were washed away by three washes with 1xPBS. Acellular CCs/CCTs (ECC/ECCTs) were collected by Unipick and kept on ice followed by storage at -80^o^C. More than 400,000 stable SILAC labeled acellular CCs/CCTs per CC/CCT proteome analysis sample were collected in > 4 years.

**Liquid chromatography tandem mass spectrometry CC/CCT proteome analyses**

The proteins in collected ECCs/CCTs with stable SILAC labeled with ^13^C_6_, ^15^N_2_-L-Lysine and ^13^C_6_, ^15^N_4_-L-Arginine were extracted and purified by SDS-gel electrophoresis. After Coomassie blue staining, the SDS-gel strip of one sample was cut into 4-5 gel slices. After in-gel digestion12.5 ng/μL trypsin, the digested peptides were extracted and enriched. The enriched peptides were used for liquid chromatography tandem mass spectrometry (LC-MS/MS) analyses as previously described^1^. The enriched peptide fractions were analyzed by liquid chromatography tandem mass spectrometry (LC-MS/MS) on an LTQ Orbitrap Velos mass spectrometer (Thermo Scientific) equipped with a Thermo Fisher Scientific nanospray source, an Agilent 1100 Series binary HPLC pump, and a Famos autosampler. Peptides were separated on a 0.125 × 180 mm fused silica microcapillary column with an in needle tip (made in-house) with a ∼5-μm i.d. The silica microcapillary column was packed with magicC18AQ C18 reverse-phase resin (5-μm particle size, 200-Å pore size; Michrom Bioresources). Separation was performed by applying a 57-min gradient from 7% to 28% acetonitrile in 0.125% formic acid. The mass spectrometer was operated with default settings: full MS [automatic gain control (AGC), 1 × 10^6^; resolution, 6 × 10^4^; m/z range, 375–1,800; maximum ion time, 1,000 ms]; MS/MS (AGC, 5 × 10^3^; maximum ion time, 120 ms; minimum signal threshold, 4 × 10^3^; dynamic exclusion time setting, 30 s; charged ions and ions for which no charge state could be determined were excluded MS/ MS selection). Triplicated independent experiments were performed.

**Database Searches, Data Filtering, Validation of Protein Detection Rate, and Proteome Analyses**.

The spectral data were searched with SEQUEST^2^ against a database containing the human protein sequence database (www.ensembl.org/index.html) together with the reversed complement. The LC-MS/MS identifications were filtered to 0.98% protein false discovery rate (FDR) and 0.1% peptide FDR. The peptide quantification and phosphorylation site localization were analyzed using in-house software and Ascore as previously described^2^.

**Cytocapsular tumorsphere and cytocapsula growth *in vitro*, immunohistochemistry staining and imaging**

Pancreas cancer Bxpc3 cells were implanted in CC/CCT culture kit (Cat. CD 0112, Celldevi) following the kit manual. At 36h, Bxpc3 cells generated cytocapsulas (CCs). Some cytocapsular oncocells performed ecellulation. Cytocapsular oncocells and ecellulated CCs were performed fixation kit and immunohistochemistry staining. At different time of 48h, 72h, 68h, 74h, 78h, 84h, 96h, 108h after cell implantation, Bxpc3 cancer cells engender cytocapsular oncocells, and grow into cytocapsular tumorspheres in different sizes with CC tightly wrapping oncocell mass or with wide cytocapsular lumens, and ecellulation of cytocapsular tumorspheres. These cytocapsular tumorspheres and ecellulated cytocapsular tumorspheres were fixed by Celldevi Inc. CC/CCT fixation kit (Celldevi, CD0201) in the 6-well plate, and then taken out and put onto slides, followed by immunohistochemistry (IHC) staining. IHC staining was performed with rabbit anti-PMCA2 polyclonal primary antibodies (1:200 dilution), mouse anti-γ-actin monoclonal primary antibodies (1:200 dilution), Goat anti-Mouse IgG (H+L) Highly Cross-Adsorbed Secondary Antibody, Alexa Fluor Plus 555 (Thermo Fisher), and Goat anti-Rabbit IgG (H+L) Highly Cross-Adsorbed Secondary Antibody, Alexa Fluor Plus 488, Thermo Fisher), and DAPI staining (1:000 dilution). Fluorescence images were taken with a Nikon 80i upright microscope with a 20× or 40× lens. All images were obtained using MetaMorph image acquisition software and were analyzed with ImageJ software.

**Histology and Immunohistochemical staining analysis**

The 9972 formalin-fixed, paraffin-embedded (FFPE) human cancer tissue specimens (4-5μm in thickness) from 9784 cancer patients, 14 human normal tissue FFPE specimens form 14 patients, and 126 human benign tumor tissue FFPE specimens from 126 patients were processed immunohistochemistry and hematoxylin and eosin (H&E) staining. Immunohistochemical fluorescence tests were performed to stain cytocapsular tubes using rabbit anti-PMCA2 polyclonal primary antibodies (1:200 dilution), mouse anti-γ-actin monoclonal primary antibodies (1:200 dilution), Goat anti-Mouse IgG (H+L) Highly Cross-Adsorbed Secondary Antibody, Alexa Fluor Plus 555 (Thermo Fisher), and Goat anti-Rabbit IgG (H+L) Highly Cross-Adsorbed Secondary Antibody, Alexa Fluor Plus 488, Thermo Fisher), and DAPI staining (1:000 dilution). Fluorescence images were taken with a Nikon 80i upright microscope with a 20× or 40× lens. All images were obtained using MetaMorph image acquisition software and were analyzed with ImageJ software.

**Time-Lapse DIC Microscopy and Videos.**

Time-lapse DIC microscopy analyses of cytocapsula elongation and cell migration were performed using a Nikon Ti motorized inverted microscope and a digital Hamamatsu ORCA-ER cooled CCD camera with a 20× lens. The time-lapse microscope was equipped with DIC, phase contrast, and epi-fluorescence optics, a Prior ProScan III mo- torized stage and shutters, a perfect focus system, and an Okolab 37 °C, 5% CO2 cage microscope incubator (Okolab). Images were taken every 30 s over the course of ∼10–36 h. All images were obtained using MetaMorph software. Tracks made by 2 h of cytocapsula elongation were obtained using MetaMorph and ImageJ software. Cytocapsula elongation velocities were also calculated using length and time measurements. Movies were prepared using the images collected via time-lapse and Meta- Morph software (15 frames/s).

**Bright filed microscope and videos**

Bright field microscope analyses of cytocapsula growth with cytocapsulasomes activities were performed using Nikon Eclipse TS2 Inverted Routine Microscope with a DS-FI3 Microscope Camera with a phase contrast 20x lens. The videos were taken using NIS-Elements software (25fps, frame per second).

**Imaging Acquisition.**

DIC and fluorescence images of fixed cells (with or without cytocapsulae) were taken with an 80i upright microscope and a digital Hamamatsu ORCA-ER cooled CCD camera with a 20× or 40× lens. The bright-field phase-contrast image was taken using a Nikon digital camera. The cytocapsula initiation ratio per high-performance field (HPF; 200×) and the number of elongated cytocapsulae per high-performance field were quantified. All images were obtained using MetaMorph image acquisition software and were analyzed with ImageJ software.

**Data collection**

Cytocapsular tubes (CTs, not sectioned, longitudinally sectioned, and cross sectioned) without degradation (3~6μm in measured diameter) were counted using a fluorescence microscope and ImageJ. The presence of CTs degrading into thick strands (1~2μm in measured diameter), thin strands (0.2~1μm in measured diameter), or the disintegration state were reported without quantification. The patients providing formalin-fixed paraffin-embedded tissue samples gave informed consent that they understood that the biopsies (needle biopsy or surgical biopsy, from US Biomax) were performed for *in vitro* research purposes only. Comparative deidentified samples of normal tissues, benign tissues, carcinoma *in situ*, cancer, paracancer, metastatic tissues with their cancer stages identified according to the tumor (T), node (N), and metastasis (M) TNM system (cancer stages: 0, I, II, III, IV) were obtained from archival materials (Tables S3-4). The cancer, paracancer and metastatic cancer tissues, in which the original cancer niches were identified by indicated cancer specific molecular markers, were identified by hospital pathology laboratories and obtained from archival material. Autopsy tissues samples have many post-life CC/CCT degradation and CCs/CCTs will not be quantified. Biopsy samples from FFPE with fresh tissues present high fidelity of CC/CCT status and CCs/CCTs are quantified and reported.

**Quantification and Statistical Analysis.**

The statistical methods used for comparisons are indicated in the relevant figure legends and in the sections below. The diameters, widths, and lengths of cytocapsulae and cytocapsular tubes were measured with MetaMorph or ImageJ. The time of individual cytocapsulae and cytocapsular tubes was counted from cytocapsula generation to acellular cytocapsula (or cytocapsular tube) decomposition. For lifetime of cytocapsular oncocell and cytocapsular tumorsphere assays, at least 20 cytocapsular oncocell or cytocapsular tumorsphere were measured per condition, and two-tailed Student’s test was used to determine statistical significance. The movie taken time was labeled as hour : minute : second (in Figs S4-5), while times after cell implantation were 96h in Fig.S4 and 108h in Figs.5. In Fig.S7, the movie taken time is just after cell implantation. The graph plots are mean ± SD. In CCT analysis in cancer types/subtypes, at least 3 samples per cancer subtype were checked (in Figs. 3A and 5, Table S2, S5). In cytocapsular tube quantitation assays, for each specimen, the number of fully intact cytocapsular tubes was counted in 5 areas (0.35mm x 0.35mm, length x width) of the sample (top, bottom, left, right, and center), and the cytocapsular tube density (CCT/mm^2^) was calculated and determined for each area. The average CCT density across the 5 sites was treated as the specimen’s overall CCT density and round up to digits. The quantitation of a humoral vessel density employs the similar method as CCT quantitation (in Figs. 12B).

**Reference:**

### 21. Everley PA, et al, Quantitative cancer proteomics: stable isotope labeling with amino acids in cell culture (SILAC) as a tool for prostate cancer research. *Mol Cell Proteomics*; 2004;3(7):729-35.

### 22. Yi TF, etal, Quantitative phosphoproteomic analysis reveals system-wide signaling pathways downstream of SDF-1/CXCR4 in breast cancer stem cells. *Proc Natl Acad Sci U S A*;2014 May 27;111(21):E2182-90.

**Figure legends:**

**Fig. S1. Detection and lifecycle of cytocapsular oncocell.**

(**A**) Representative bright filed (BF) and immunohistochemistry (IHC) microscope images of cytocapsular tube (CCT) in CC/CCT culture kit matrix *in vitro*. There is a single Bxpc3 cancer cell in migration in the CCT. CCT membrane edges (orange arrows) are shown. (**B**) Representative immunohistochemistry microscope image of cytocapsular tumorsphere (CT, purple arrows) in CC/CCT culture kit matrix *in vitro*. Cytocapsula (CC, white arrow), ecellulated CC (ECC, orange arrows) and ECC with open holes (white asterisk) are shown. (**C**) Representative immunohistochemistry microscope image of human normal, benign and cancer tissues with CC/CCT detection by anti-PMCA2 antibodies and anti-γ-actin antibodies. Terminal duct (TD), cytocapsular tube (CCT, white arrows), CCT strand (CTS, orange arrows) are shown. (**D**) Schematic diagram of lifecycle of cytocapsular oncocell: transformed (cancerous) cells experience devolution and perform cytocapsulasome-driven cytocapsula formation. Cytocapsulas wrap oncocells inside and form cytocapsular oncocells. (1) cytocapsular oncocells proliferate and grow into cytocapsular tumors. (2) Cytocapsular oncocells proliferate and cytocapsulas elongate and develop into CCTs. (3) Ecellulation of cytocapsular oncocells produces acellular cytocapsulas and acytocapsular oncocells. Acellular cytocapsulas and CCTs perform autodegradation and decomposition and disappearance.

**Fig. S2. Detection of CCTs by IHC fluorescence staining with anti-PMCA2 antibodies.**

(**A-B**) Representative image of H&E staining (A) and IHC fluorescence staining with anti-PMCA2 antibodies (B) of two continuously sectioned and neighboring colon carcinoma tissue specimens. Panel 1 is the microscope image of a whole colon carcinoma core. The framed area in panel 1 is enlarged and shown in panel 2. The framed area in panel 2 is enlarged and shown in panel 3. (**C-D**) Representative image of H&E staining (C) and IHC fluorescence staining with anti-PMCA2 antibodies (D) of two continuously sectioned and neighboring lung carcinoma tissue specimens. CCTs in colon and lung carcinoma tissues are invisible in H&E technologies (panels A and C), but are clearly shown in images with IHC fluorescence staining with anti-PMCA2 antibodies (panels B and D). Cytocapsular tube (CCT, white arrows), and CCT strand (CTS, orange arrows).

**Fig. S3. Detection of CCTs, CCT formation and cytocapsular tumor lifecycle.**

(**A**) Representative image of H&E staining (1), IHC staining with antibodies recognizing breast cancer molecular markers ER (2), PR (3), and HER (4), and IHC staining with anti-PMCA2 antibodies (5) of 5 pieces of continuously sectioned breast cancer tissue specimens. The framed areas in the same site of each tissue specimen show that CCTs presented in panel 5 are invisible in the panels 1-4. (**B**) Representative image of H&E staining (1), IHC staining with antibodies recognizing colon cancer molecular markers MSH-2 (2), and IHC staining with anti-PMCA2 antibodies (3) of 3 pieces of continuously sectioned colon cancer tissue specimens. The large quantities of CCTs presented in panel 3 are undetectable in panels 1 and 2. (**C**) Representative bright field (BF) images and fluorescence microscope images of the same imaged area show that a long, straight, and stretched CCT links two cytocapsular tumorspheres (CTs) with open ends in both connection sites. Cytocapsular tube (CCT, white arrows), cytocapsulas (CC1 and CC2) of the two CTs, and open end (cyan arrows) of CCT are shown. (**D**) Schematic diagram of lifecycle of cytocapsular tumor: (1) Generation of cytocapsular oncocells; (2) cytocapsular oncocells proliferate and grow into prophase cytocapsular tumor (PCT) without CCTs. Incytocapsular oncocells may experience devolution and generate independent CCs in the PCT cytocapsular lumen. (3) PCT develops into CT with CCTs. CCTs provide physical membrane-enclosed freeways for incytocapsular oncocell metastasis. (4) CCs of cytocapsular tumors and CCTs degrade, and form acytocapsular oncocell masses of NT. (5) Some acytocapsular oncocells experience devolution in the stressful microenvironments in NTs, and generate cytocapsular oncocells, which engender CCTs, and form Acytocapsular oncocell Mass-CT/CCT Complex (AMCC). Acytocapsular oncocells around CCTs invade into CCTs via alloentry and proceed cancer metastasis.

**Fig. S4. Cytocapsulasomes drives cytocapsula growth.**

Real-time analysis of cytocapsula growth driven by cytocapsulasomes with a bright field phase contrast microscope. The images are taken from Movie S1. Cytocapsulasome activity procedures: (1) cytocapsulasome (CS) are generated and released by oncocells, and attach to the outside cytoplasm membrane (cyan arrows); (2) CSs spontaneously are detached from oncocell surface (panels 5 and 6, orange arrows); (3) detached CSs randomly move in the cytocapsular lumen fluids (yellow arrows in panel 4, 8, 9, 10, 11); (4) CSs reach the inner side of cytocapsular membrane of CT, and contact and integrate into cytocapsular (CC) membrane of CT, and increase CC membrane size in area (red arrows, panels 8 and 9). In cytocapsular tumors with cytocapsula tightly wrapped oncocell mass surfaces, the contact and integration of cytocapsulasomes into CC membranes will be faster and more efficient without random and long movement journey in the cytocapsular lumen fluids. Incytocapsular tumorsphere (ICT), cytocapsula (CC, white arrows), cytocapsular membrane (CCM, green arrow), cytocapsular spike (purple arrows), and cytocapsulasome (CS; yellow arrow, CS in random movement in cytocapsular lumen fluid; orange arrow, CS detaching from oncocell surface; and red arrows, CS reach and fuse into CC membrane; cyan arrow: CS attached on oncocell surface) are shown.

**Fig. S5. Large quantities of cytocapsulasomes promote cytocapsular growth.**

(**A**) Real-time analysis of cytocapsula growth driven by cytocapsulasomes with a bright field phase contrast microscope. The images are taken from Movie S2. Incytocapsular tumorsphere (ICT), cytocapsula (CC, black arrows), cytocapsula membrane (CCM, orange arrow), cytocapsulasome (CS, red and green arrows), ecellulated cytocapsula (ECC), and cytocapsular tube (CCT, in the big CC lumen) are shown. (**B**) Quantitation of cytocapsulasomes in cytocapsular tumorspheres.

**Fig. S6. Characterization of cytocapsulasome and its lifecycle.**

(**A**) Representative image of cytocapsulasomes (CSs, white arrows) in gastric cancer cells during the generation of cytocapsular oncocells *in vivo*. There are many CSs in the cytoplasm. (**B**) Quantitation of cytocapsulasomes in gastric and breast cancer cells with cytocapsulasomes *in vivo*. (**C**) Comparative differences and similarities between cytocapsulasome and other 4 extracellular vesicles. (**D**) Schematic diagram of cytocapsulasome lifecycle: (1) Generation of CS in the cytoplasm, (2) Release of CS onto the outside of cytoplasm membrane, and attach to the cell membrane surface, (3) Multiple CSs on the cell membrane contact and integrate into cytocapsular membrane fragments, (4) Cytocapsular membrane fragment grow up with more CS integration, (5) Cytocapsular membranes envelope the whole single cell and generate cytocapsular oncocells, and isolate them from the ECM, (6) CSs reach and integrate into cytocapsular membranes and increase cytocapsular membrane areas, (7) Cytocapsulas grow up and generate enlarged cytocapsulas or elongate and develop into cytocapsular tubes.

**Fig. S7. Cytocapsular tube generation and elongation, and cell migration in CCTs *in vitro* and *in vivo***

(**A**) Real-time analysis of cytocapsular tube initiation, generation and elongation. The images are taken from Movie S3. Cytocapsular tube (CCT, red arrows), cell in CCT (white arrows), bleb (cyan arrows), cytocapsular spike (CSP, yellow arrows) are shown. (**B**) Quantitation of cytocapsular tube elongation speed in the CC/CCT culture kit matrix *in vitro*. (**C**) Representative image of initial cytocapsular tube (IC, white arrows) in breast cancer tissues *in vivo*. (**D**) Schematic diagram of cytocapsular tube elongation: (1) Incytocapsular oncocell generate and release cytocapsulasomes and drive cytocapsular membrane area increase, (2)cytocapsular oncocells generate many blebs in all directions, sense microenvironments, choose and decide the motility directions, (3) cytocapsulasomes continuously drive cytocapsulas (CC) membrane increase in areas with many blebs in the CC lumen, (4) cytocapsular oncocells move forward, and elongate CCT length, and generate long CCTs. (**E**) Representative image of cell migration in CCT *in vitro*. (**F**) Real-time analysis of incytocapsular oncocell migration in CCTs. The images are taken from Movie S4. Cytocapsular tube (CCT, red arrows), incytocapsular oncocell in migration (white arrows), migration direction (pink arrows), reversed migration direction (yellow arrows), cell with lamellipodia at the leading edge (red asterisk), cell in transition of migration direction in CCT (blue asterisk) are shown. (**G**) Quantitation of Bxpc3 pancreas cancer cell migration in CCT *in vitro*. (H) Representative images of colon cancer cells in migration in colon CCTs. Colon cancer incytocapsular oncocells in CCTs are in thin, long and spindle-shaped morphologies. Cytocapsular tube (CCT, red arrows), and colon oncocells in migration in CCT (white arrows) are shown. (I) A schematic diagram of cytocapsular tube tumor.

**Fig. S8. Integrated cytocapsular oncocell, cytocapsular tumorspheres, CCT networks, and cytocapsular tumorsphere network systems *in vitro*.**

(**A**) Representative image of primary and secondary cytocapsular tumorspheres integrated cytocapsular tumorsphere network systems (CTNSs) *in vitro*. Firstly, primary cytocapsular tumorspheres interconnect by CCT networks. Disseminated incytocapsular oncocells gather in the CCT network nodes and grow into secondary cytocapsular tumorspheres. Secondary cytocapsular tumorspheres interconnect primary cytocapsular tumorspheres via CCTs and generate a combined primary and secondary cytocapsular tumorsphere network system in the CC/CCT culture kit (6-well plate). (**B**) Representative image of cytocapsular tumorsphere interconnection by CCTs with open-ends in both sides of CCT, and incytocapsular oncocell migration in CCTs and CTNSs. (**C**) Representative image of membrane-sheltered cytocapsular oncocells, cytocapsular tumorspheres, CCT networks, and integrated CTNSs. Cytocapsular tube (CCT, white arrows), cytocapsula (CC, yellow arrows), open-end of both sides of CCTs (cyan arrows) , and incytocapsular oncocell migration in CCTs (red arrows) are shown.

**Fig. S9. Lifecycle of primary cytocapsular tumor network systems *in vivo*.**

(**A**) Representative image of dense prophase cytocapsular tumor (PCT) groups in primary invasive ductal breast carcinoma. Degradation of acellular cytocapsula (CC) leads to cloud-like CC strand masses (orange arrows). (**B**) Quantitation of PCT density in breast, colon and prostate cancers. (**C**) Highly dense cytocapsular tumor (CT) groups in primary invasive ductal breast carcinoma. The framed area is enlarged and shown in Fig. 4A. (**D**) Quantitation of CT density in breast, colon and prostate cancers. (**E**) Representative image of CCT networks in primary CTNSs in primary colon cancer. (**F**) Early nuperphase tumor. The CT cytocapsula is degraded with CC fragments (orange arrow) remain. CCTs degrade into CCT strands (CTSs). (**G**) Late stage of nuperphase tumor. There are dense acytocapsular oncocells without CCTs. (**H**) Representative image of acytocapsular oncocell mass-CC/CCT complex (AMCC) in primary breast cancer. many new CCTs are regenerated by some acytocapsular oncocells. (**I**) CCT degradation in AMCC. (**J**) Many acytocapsular oncocells invade into CCTs via alloentry, metastasize and leave away. The local oncocell density is very low. Many CCTs are in degradation into strands. (**K**) Severe CCT degradation with many CCT fragments and strands. (**L**) CCT degradation and oncocell apoptosis in cytocapsular tumor lumens.

**Fig. S10. CCTs are broadly present in native tissue adjacent to tumors (NAT).**

(**A**) Representative image of NAT in primary breast carcinoma with large quantities of CCT networks in high density. (**B**) Representative image of NAT in bone marrow of primary plasma cell myeloma with CCT bunches coexist with immune cells. (**C**) Representative image of NAT of trabecular bone in primary plasma cell myeloma. CCTs invade into hard tissues of trabecular bone. (**D**) Quantitation of CCT density in NAT of 14 kinds of tissues.

**Fig. S11. Cytocapsular oncocell metastasis in primary CTNSs in primary cancer niche.**

(**A**) Representative image of CCTs in loose soft tissue in lung cancer. (**B-C**) CCTs in compact soft tissues in thyroid (B) and oral (C) cancers. (**D**) CCT superstructures in pancreas cancer. (**E**) CCT networks composed by straight CCTs in primary breast cancer. (**F**) CCT networks composed by curled and coiled CCTs in primary breast cancer. (**G**) Massive curled CCT networks invade into compact acytocapsular oncocell masses and form AMCC. (**H**) Dense CCT network masses invade through compact primary prostate cancer tissues in AMCC. (**I**) Representative image of CCT bunches in primary CTNSs. (**J**) Representative images of CCT masses in primary CTNSs and most acytocapsular oncocells left via CCTs. (**K**) Representative image of highly curled and coiled CCTs in primary CTNSs in primary pancreas cancer with many acytocapsular oncocells left via CCTs. (**L**) Representative image of complex CCT superstructures in primary CTNSs.

**Fig. S12. Cytocapsular tube networks dominate cancer metastasis *in vivo*.**

(**A**) Representative image of CCTs and blood vessel in primary prostate cancer tissues. (**B**) Quantitative comparison of cytocapsular tubes and humoral vessels in primary cancer niche, normal tissue adjacent to the tumor (NAT), and secondary cancer niche. (**C**) Representative image of micro blood vessels wrapped by CCTs, CCT invasion into blood vessels, and release of incytocapsular oncocells into the blood as resources of circulating tumor cells(CTCs). Framed area is enlarged and shown in (D). (**D**) Enlarged area from (C). The incytocapsular oncocell (ICO, yellow arrow) entering blood is shown. (**E**) Representative image of late cancer stage with many red blood cells distributed in tissues caused by CCT invasion-damaged blood vessels. (**F**) Schematic diagram of CCT invasion, CTC resource, micro blood vessel damage, and red blood cell distribution in tissues beyond blood vessels.

**Fig. S13. Lifecycle of secondary cytocapsular tumor network systems *in vivo*.**

(**A**) Representative images of metastatic breast cancer CCTs wrapping and invading lymph nodes. (**B**) Metastatic cervix squamous cell carcinoma CCT networks invade into and distributed inside most lymph node areas. (**C**) Representative images of secondary breast CTNSs with high cytocapsular tumor density in lymph node. Breast CCTs invade into lymph node, incytocapsular oncocells proliferate and grow into cytocapsular tumors. Large quantities of secondary breast cytocapsular tumors in lymph node interconnect by CCT networks and form dense CTNSs in lymph node. (**D**) Quantitation analysis of secondary breast CTs in bladder, liver and lymph node. (**E**) CTs in secondary CTNSs grow into bigger CTs in lymph node. (**F**) Degradation of CCs of secondary CTs, CCT degradation, and acytocapsular ovary oncocell uncontrolled proliferation led to big acytocapsular ovary oncocell masses in secondary CTNSs in omentum. (**G**) Some acytocapsular ovary oncocells regenerate CCTs, and form AMCC in secondary CTNSs in omentum. (**H**) Representative image of secondary cervix CTNSs in lymph node with many cervix CCTs in lymph node. (**I**) Representative image of a whole tissue core of secondary rectum CTNSs in mesentery with high CCT density. Framed area is enlarged and shown in (J). (**J**) Enlarged area from (I) to show the high CCT density. (**K**) Secondary colon CTNSs with severe CCT degradation in liver. (**L**) Secondary hepatocellular carcinoma CTNSs with severe CCT degradation, severe blood vessel damage, and severe red blood cell leak, major acytocapsular oncocell metastasis via CCTs, and low cell density in the local cerebrum.

**Fig. S14. Schematic diagram of an atlas of cytocapsular oncocell evolution lifecycle *in vivo*.**

A simplified atlas of cytocapsular oncocell evolution lifecycle *in vivo* includes 10 steps: (1) Normal cell transformation generates abnormal acytocapsular oncocell caused by accumulated gene mutation and chemical and physical stimuli from extracellular microenvironments. (2) Uncontrolled proliferation of acytocapsular oncocell engender acytocapsular oncocell mass, resulting in local blood supply insufficiency, nutrient deprivation, hypoxia, hypoxic metabolism, accumulated lactic acids, low p*H*, decreased intracellular space (<5%, or even 0), insufficient waste metabolic molecule effusion, elevated waste metabolic molecules, increased viscosity, and stressful microenvironments. The stressful microenvironments lead to apoptosis of some acytocapsular oncocells. Some acytocapsular oncocells survive via devolution: generation of cytocapsulas enclosing the cell and isolating the cell from stressful microenvironments. The additional extracellular protective cytocapsula of cytocapsular oncocell advance to survive under stressful microenvironments. (3) Incytocapsular oncocell proliferate in cytocapsular lumen and generate prophase cytocapsular tumor (PCT) and cytocapsular tumor (CT). (4) CTs in the primary niche generate CCTs, CCT networks, and acytocapsular oncocell mass-CC/CCT complex (AMCC). (5) All CTs in primary niches interconnect by CCT networks and form primary cytocapsular tumor network systems (CTNSs). (6) Primary CCT networks expand and invade into neighboring and far distance tissues and organs. (7) CCT branching morphogenesis, new CCT network formation, and new CT formation and growth in secondary niches. (8) All CTs in the secondary niche interconnect with CCT networks and form secondary CTNSs. AMCC formation in secondary niches. (9) Primary and secondary CTNSs have existed interconnections with CCT metastatic CCT networks, and form integrated primary and secondary CTNSs. (10) A series of activities and responses of CTs, CCTs, CCT networks and CTNSs under various conditions shape the dynamic integrated CTNSs. The expansion and invasion of CCTs, CTs, CTNSs and AMCCs in normal tissues and organs lead to normal cell apoptosis, normal tissue biological function failure and structure damage.
