## Supplementary material for "Malignant tumors engender and employ cancer-specific organelles for self-protection and metastasis": Supple figures

### Fig. S1

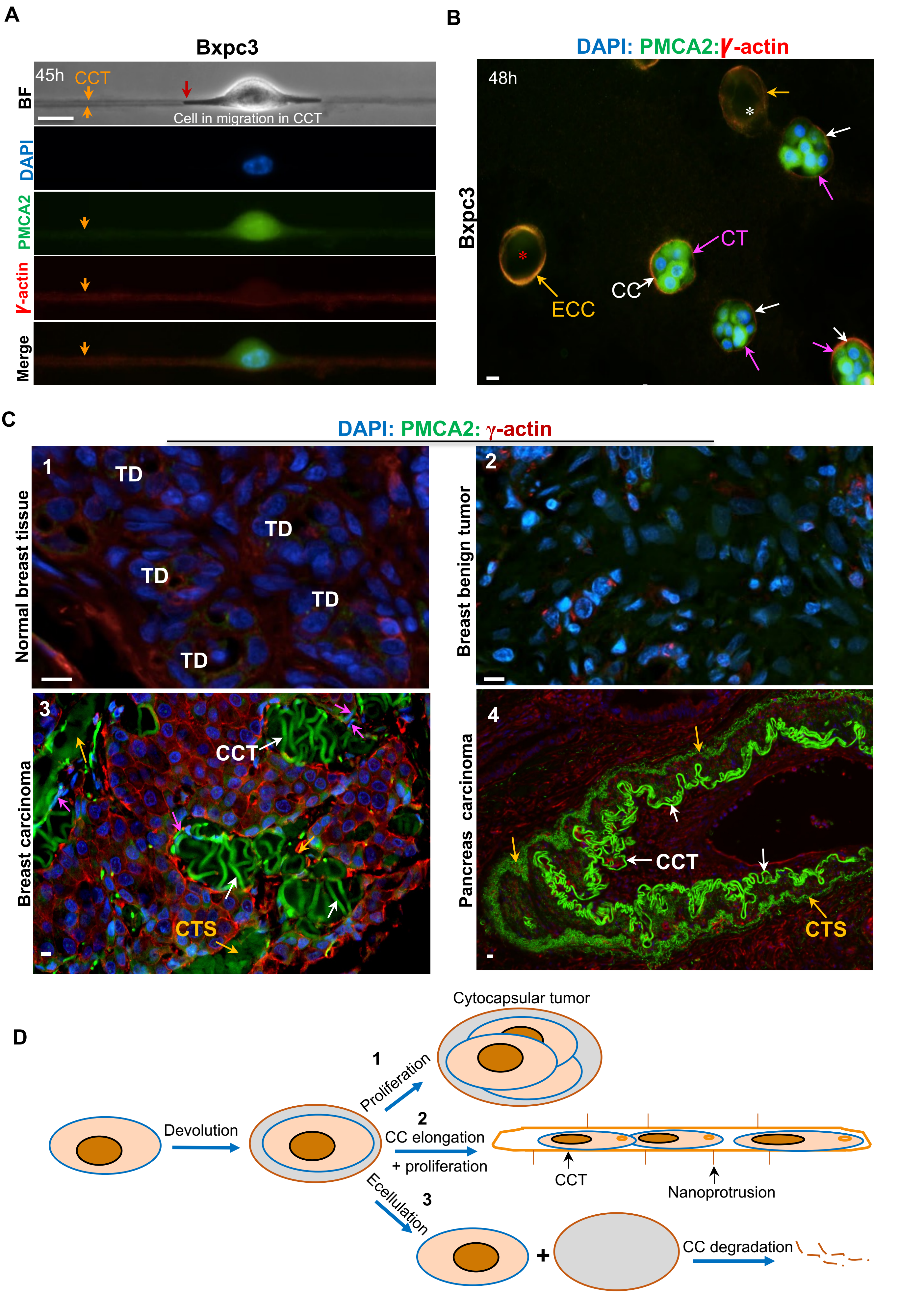

A
Bxpc3
45h
CCT
Cell in migration in CCT
BF
DAPI
PMCA2
γ-actin
Merge
B
DAPI: PMCA2:γ-actin
48h
Bxpc3
CT
CC
ECC
C
DAPI: PMCA2: γ-actin
1
TD
Normal breast tissue
2
Breast benign tumor
3
CCT
CTS
Breast carcinoma
4
CCT
CTS
Pancreas carcinoma
D
Cytocapsular tumor
1
Proliferation
Devolution
2
CC elongation
+ proliferation
CCT
Nanoprotrusion
Ecellulation
3
CC degradation

Fig. S2

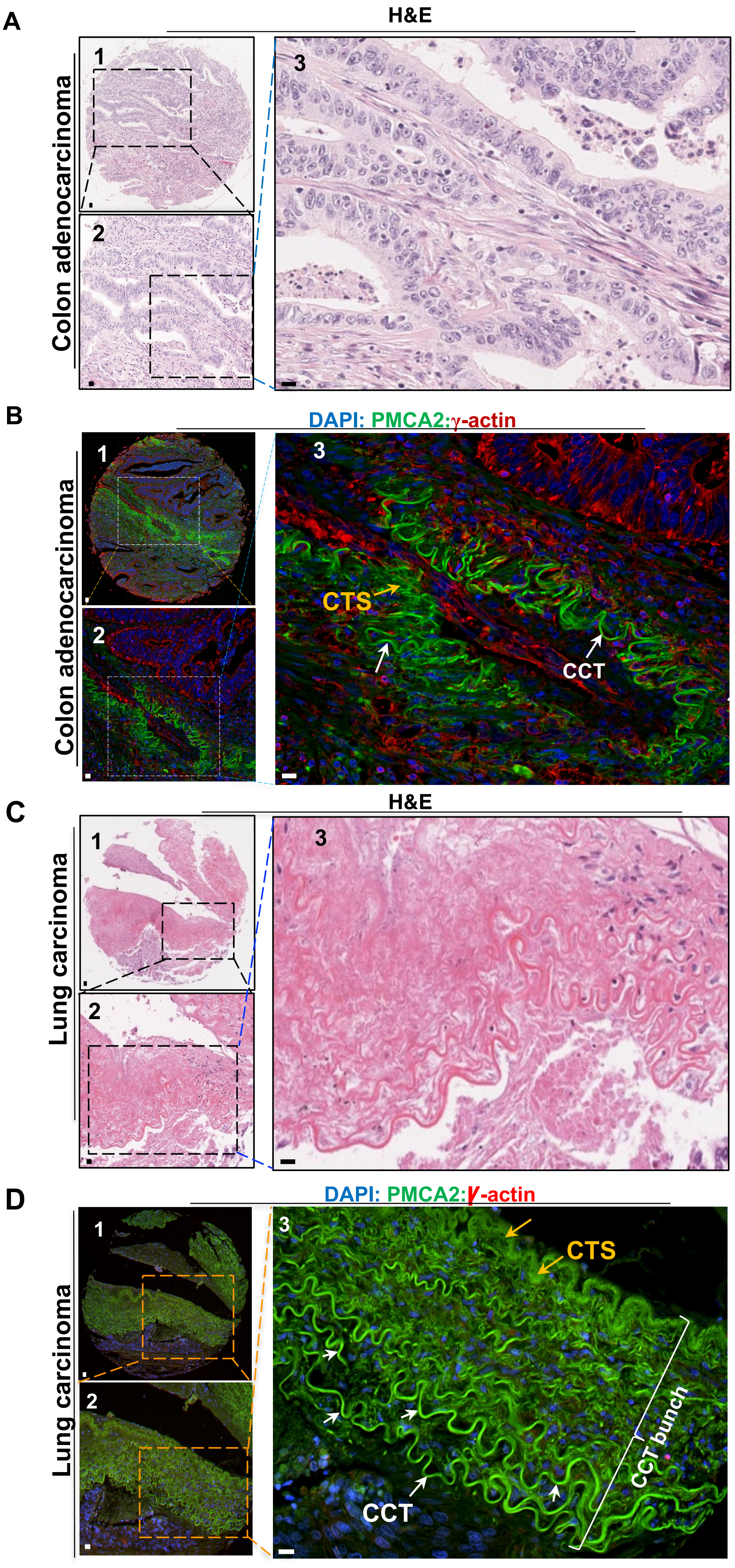
A
H&E
Colon adenocarcinoma
1
2
3
B
DAPI: PMCA2:γ-actin
Colon adenocarcinoma
CTS
CCT
C
H&E
Lung carcinoma
D
DAPI: PMCA2:γ-actin
Lung carcinoma
CTS
CCT
CCT bunch

Fig.S3

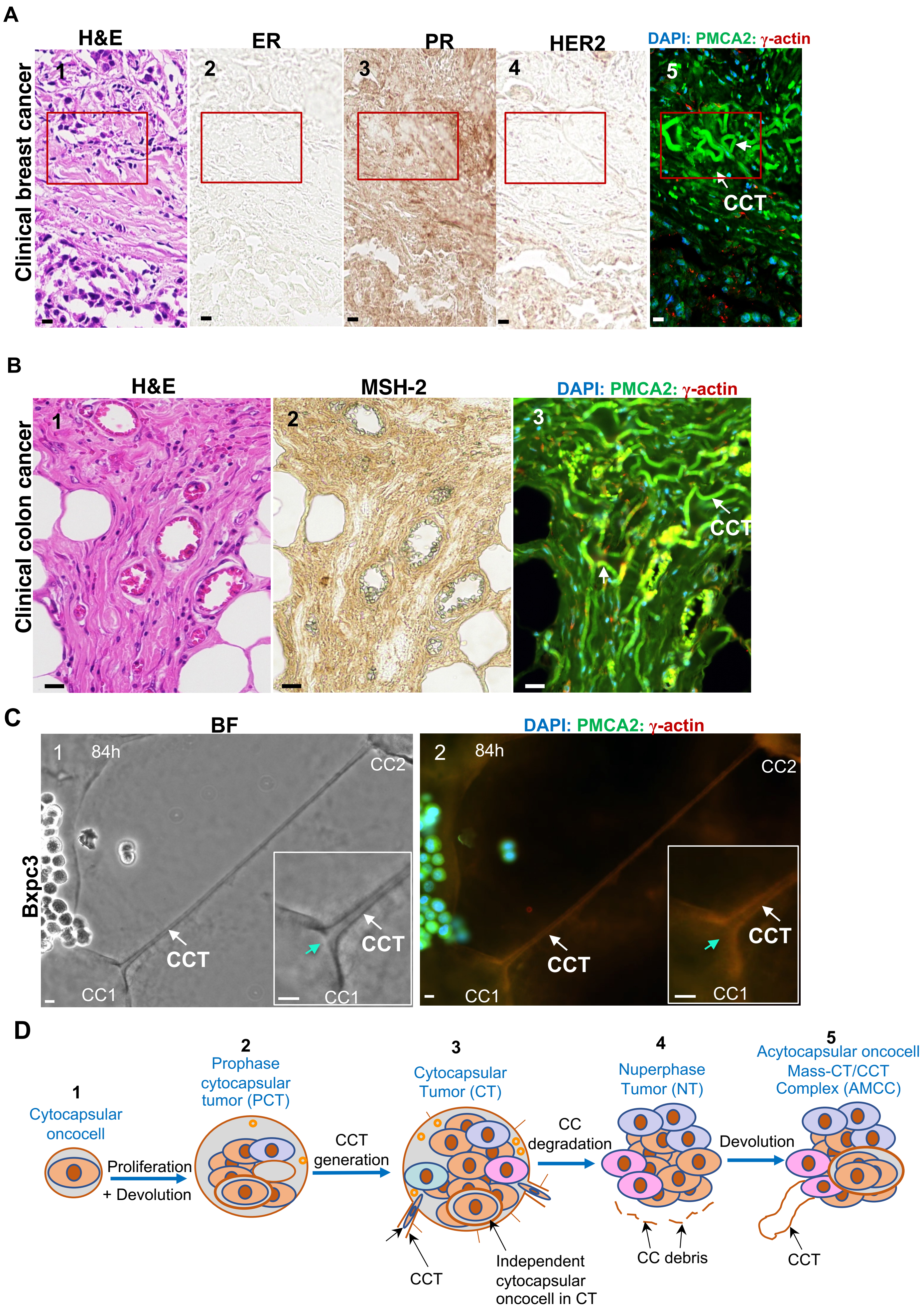
A
H&E
ER
PR
HER2
DAPI: PMCA2: γ-actin
Clinical breast cancer
1
2
3
4
5
CCT
B
H&E
MSH-2
DAPI: PMCA2: γ-actin
Clinical colon cancer
1
2
3
CCT
C
BF
DAPI: PMCA2: γ-actin
Bxpc3
1 84h
2 84h
CC2
CC1
CCT
D
1
Cytocapsular oncocell
Proliferation
+ Devolution
2
Prophase cytocapsular tumor (PCT)
CCT generation
3
Cytocapsular Tumor (CT)
CCT
Independent cytocapsular oncocell in CT
CC degradation
4
Nuperphase Tumor (NT)
CC debris
Devolution
5
Acytocapsular oncocell Mass-CT/CCT Complex (AMCC)
CCT

Fig.S4

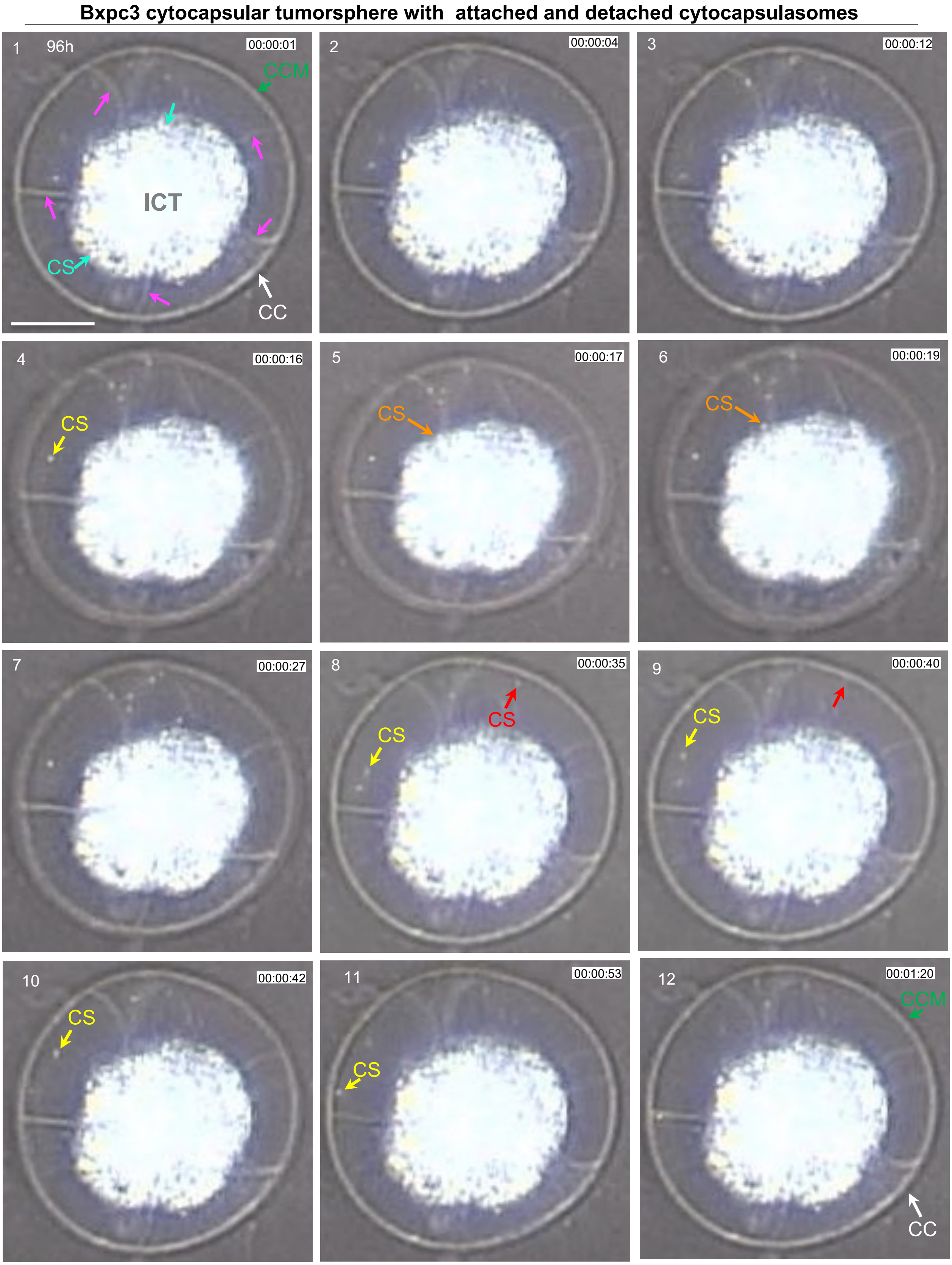
Bxpc3 cytocapsular tumorsphere with attached and detached cytocapsulasomes
1
96h
00:00:01
CCM
ICT
CS
CC
2
00:00:04
3
00:00:12
4
00:00:16
CS
5
00:00:17
CS
6
00:00:19
CS
7
00:00:27
8
00:00:35
CS
CS
9
00:00:40
CS
10
00:00:42
CS
11
00:00:53
CS
12
00:01:20
CCM
CC

**Fig.S5**

**A**

**Bxpc3 cytocapsular tumorsphere with large quantity of cytocapsulasomes in movement**

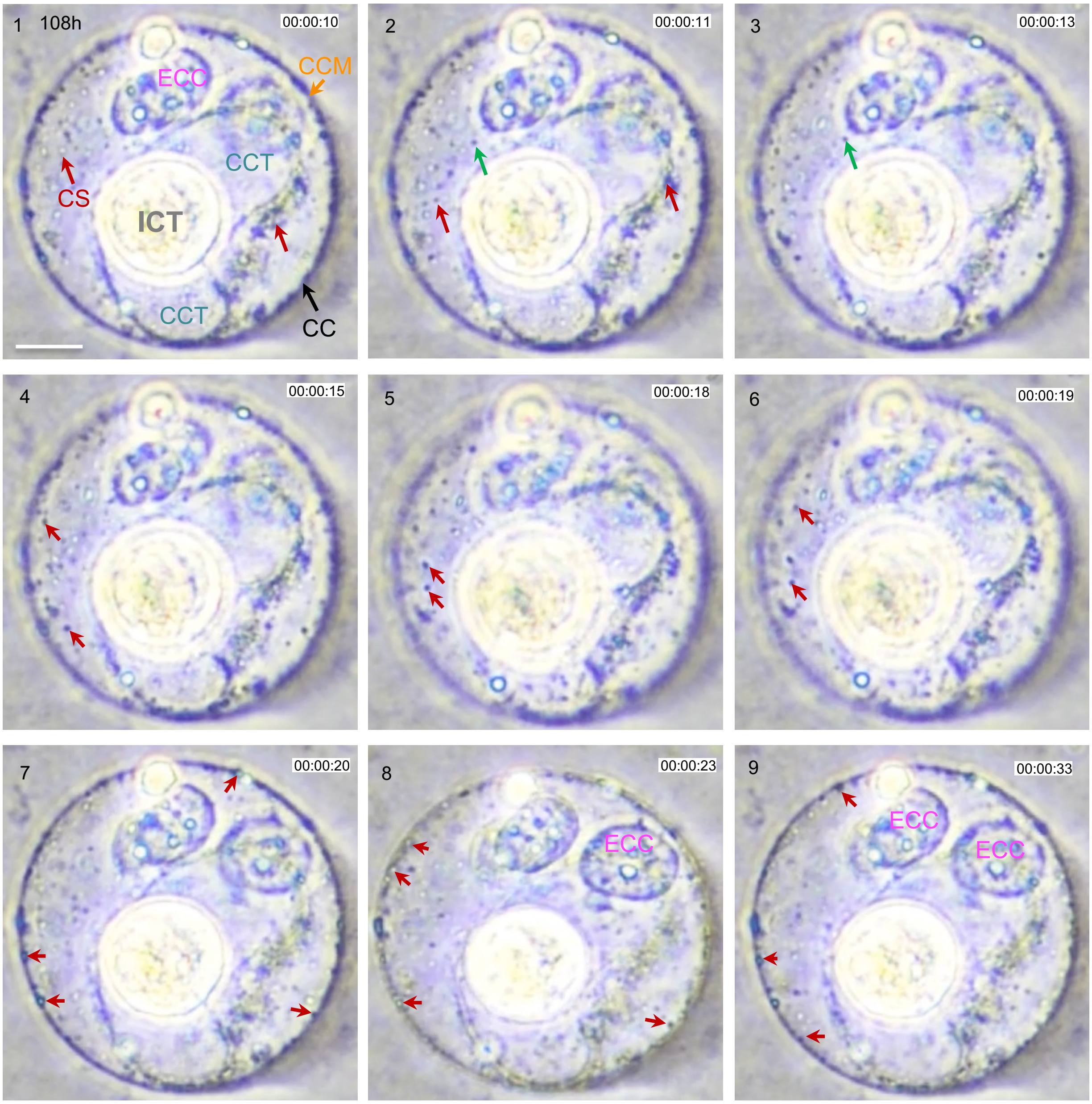


**B**

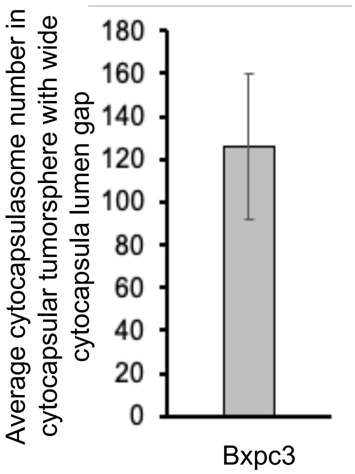

**Fig. S6**

**A**

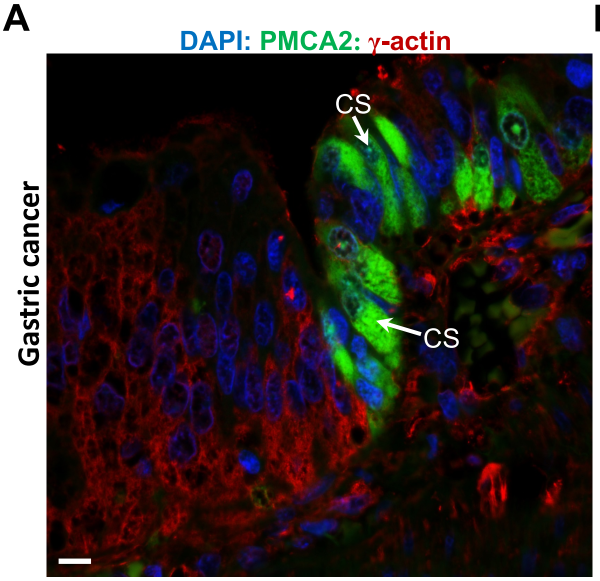


**B**

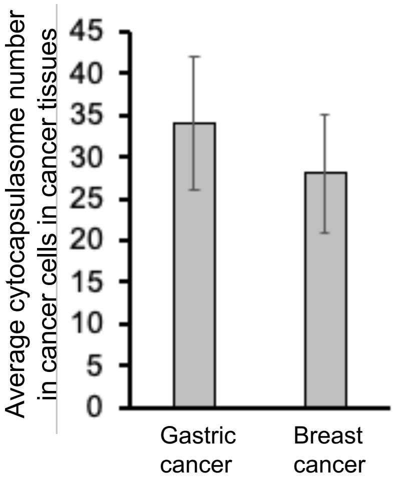


**C**

**Comparative differences and similarities between cytocapsulasome and other 4 extracellular vesicles.**

| Feature | Cytocapsulasome | Exosome | Apoptotic body | Microvesicle | Large oncosome |
|---|---|---|---|---|---|
| Topology | Sphere | Sphere | Sphere or irregular morphology | Sphere or irregular morphology | Sphere or irregular morphology |
| Membrane enclosed vesicles | + | + | + | + | + |
| If merge to form cytocapsula | + | - | - | - | - |
| If merge to from cytocapsular tube | + | - | - | - | - |
| Homogeneous or heterogeneous in size | Heterogeneous | Homogeneous | Heterogeneous | Heterogeneous | Heterogeneous |
| Size in diameter/width | 40-500nm | 30-100 nm | 1–5 μm | 100-1000 nm | 1-10 μm |
| Localization | In CC/CCT lumen | Outside cell | | Outside cell | Outside cell |
| Cell type that generates vesicle | Cytocapsular oncocell | Cell | Cell | Cell | Cell |
| Marker proteins | PMCA2 | Membrane impermeable (PI negative) CD63, TSG101, Alix | Membrane permeable (PI positive) Annexin V | | EGFRvIII |
| Destinations | Inner side of CC and CCT membranes | Other cells | Intercellular spaces | Other cells | Other cells |
| Biological functions | CC and CCT building blocks， ECM degradation, and other functions | Intercellular communications | Cellular apoptosis | Intercellular communications | Transferring tumor-promoting material |

**D**

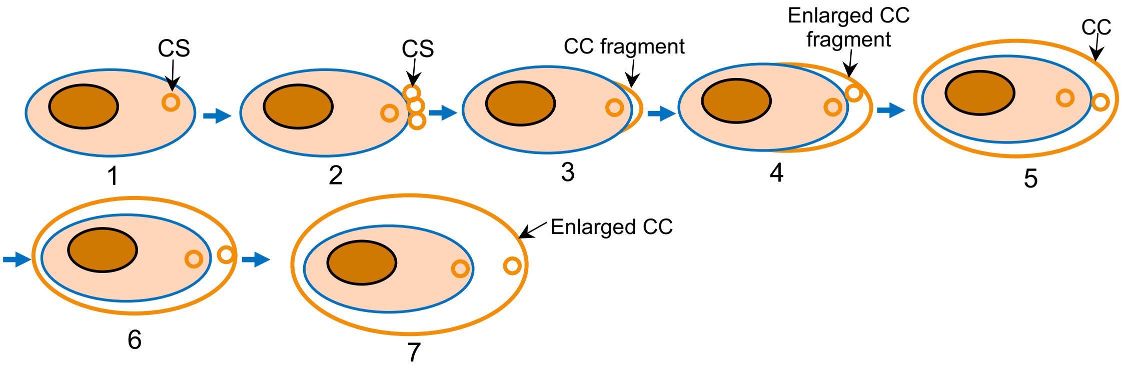

### Fig. S7

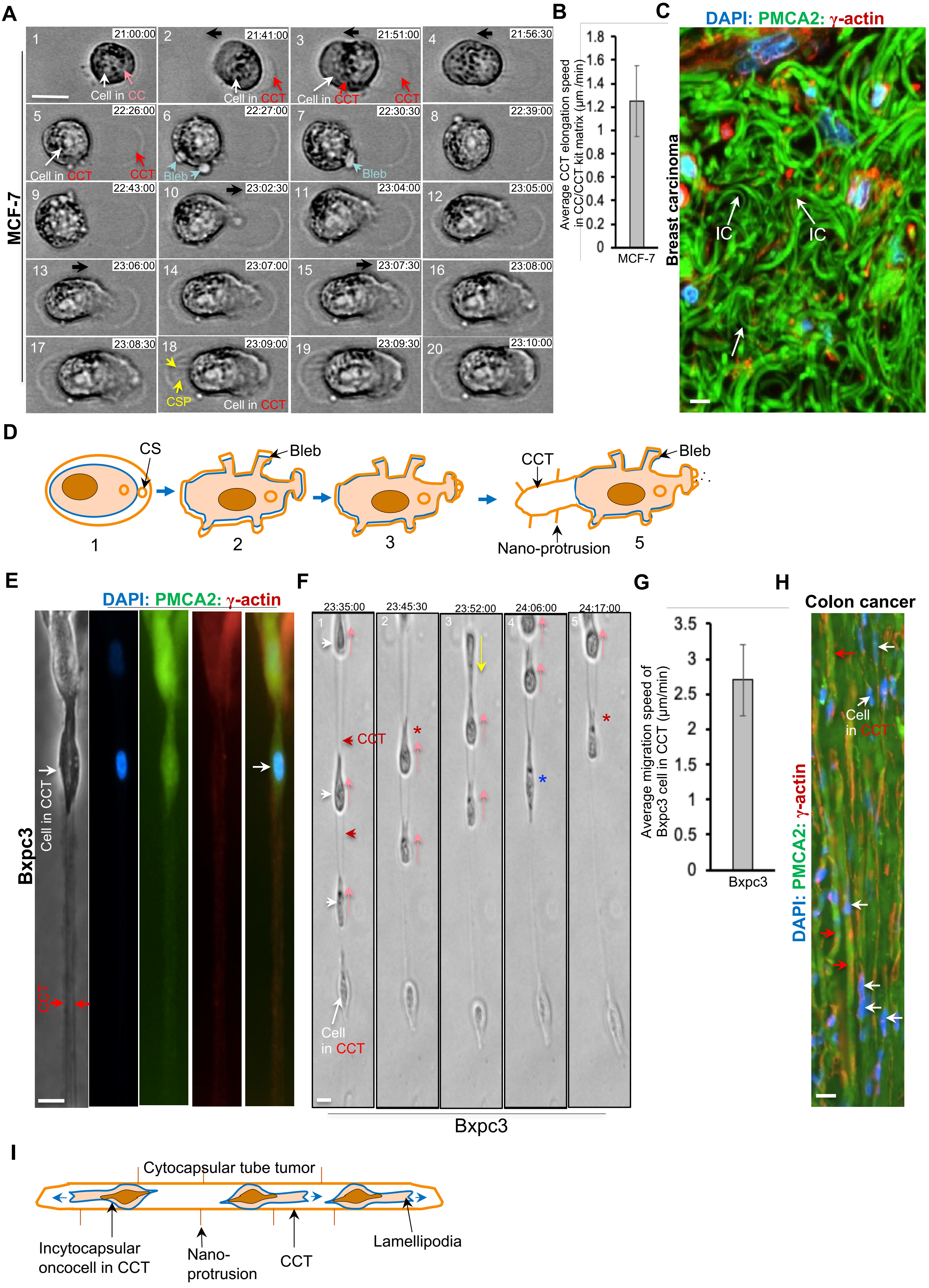
A
MCF-7
Cell in CC
Cell in CCT
CCT
Bleb
CSP
B
Average CCT elongation speed in CC/CCT kit matrix (μm /min)
MCF-7
C
DAPI: PMCA2: γ-actin
Breast carcinoma
IC
D
CS
Bleb
CCT
Nano-protrusion
E
DAPI: PMCA2: γ-actin
Bxpc3
Cell in CCT
CCT
F
23:35:00
23:45:30
23:52:00
24:06:00
24:17:00
Cell in CCT
Bxpc3
G
Average migration speed of Bxpc3 cell in CCT (μm/min)
Bxpc3
H
Colon cancer
Cell in CCT
DAPI: PMCA2: γ-actin
I
Cytocapsular tube tumor
Incytocapsular oncocell in CCT
Nano-protrusion
CCT
Lamellipodia

Fig. S8

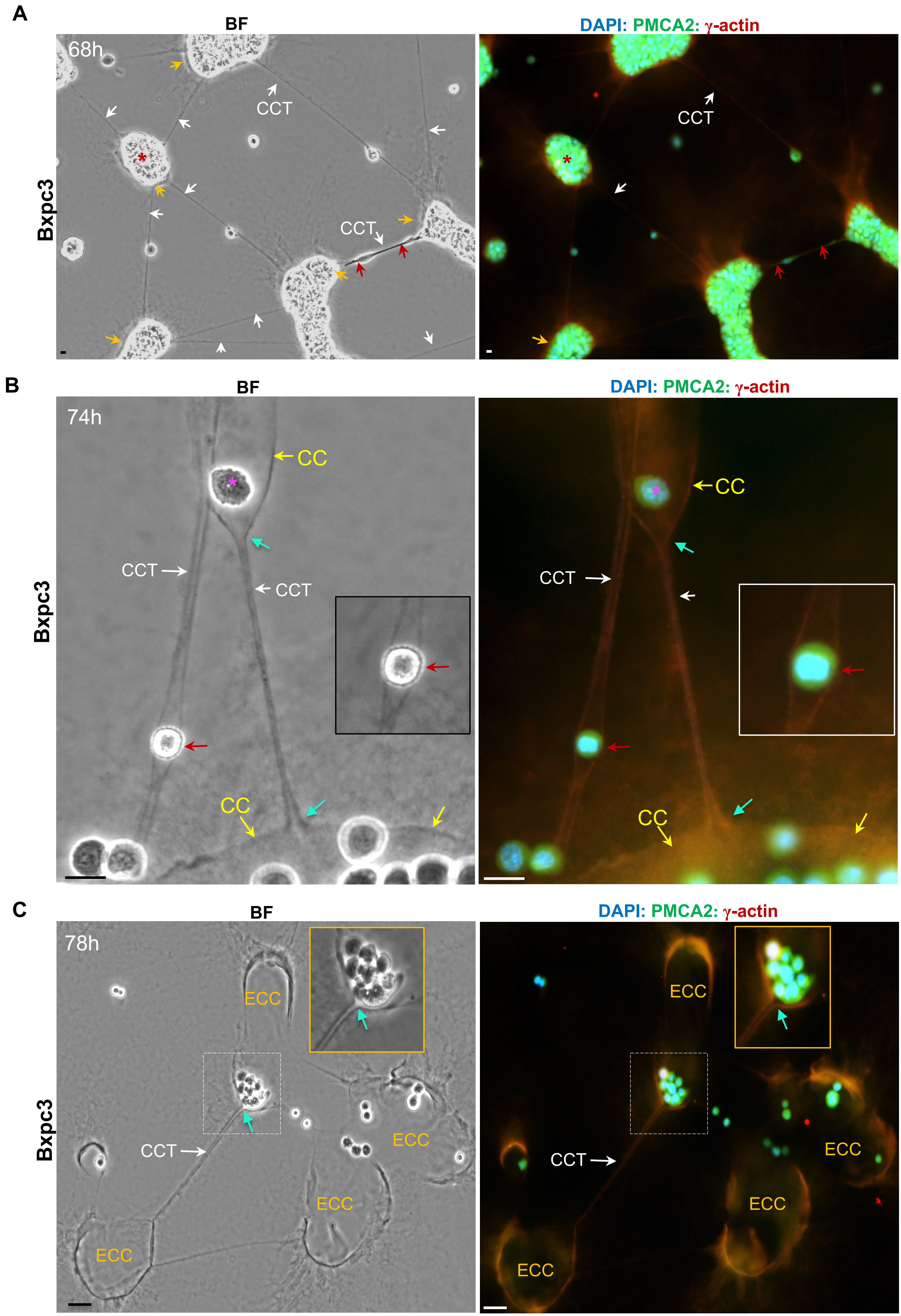

A
BF
DAPI: PMCA2: γ-actin
68h
CCT
CCT
CCT
Bxpc3
B
BF
DAPI: PMCA2: γ-actin
74h
CC
CC
CCT
CCT
CCT
CC
CC
Bxpc3
C
BF
DAPI: PMCA2: γ-actin
78h
ECC
ECC
ECC
ECC
CCT
ECC
ECC
ECC
ECC
CCT
Bxpc3

Fig.S9

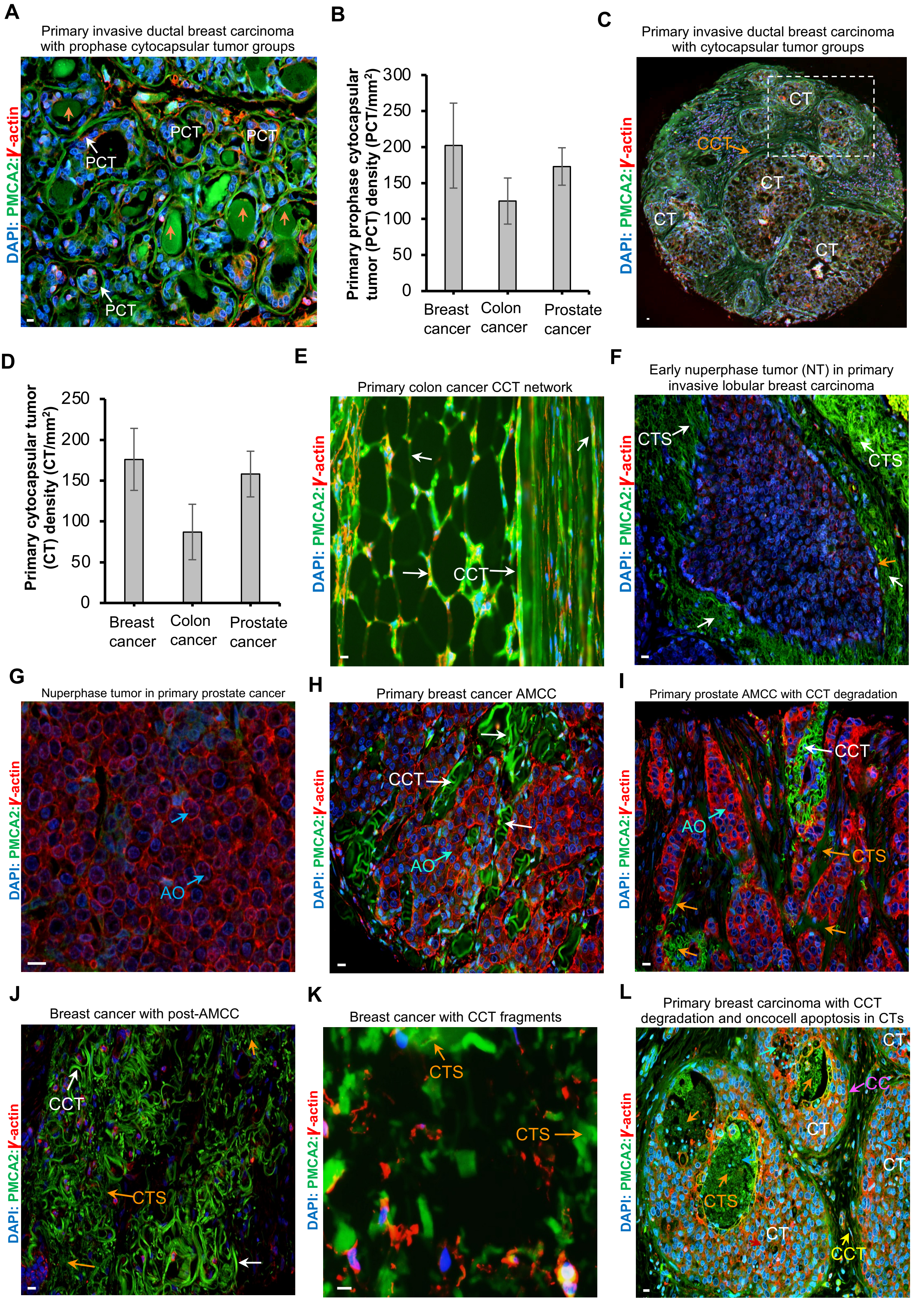
A
Primary invasive ductal breast carcinoma with prophase cytocapsular tumor groups
DAPI: PMCA2:γ-actin
PCT
PCT
PCT
PCT
B
Primary prophase cytocapsular tumor (PCT) density (PCT/mm²)
300
250
200
150
100
50
0
Breast cancer
Colon cancer
Prostate cancer
C
Primary invasive ductal breast carcinoma with cytocapsular tumor groups
DAPI: PMCA2:γ-actin
CT
CCT
CT
CT
CT
D
Primary cytocapsular tumor (CT) density (CT/mm²)
250
200
150
100
50
0
Breast cancer
Colon cancer
Prostate cancer
E
Primary colon cancer CCT network
DAPI: PMCA2:γ-actin
CCT
F
Early nuperphase tumor (NT) in primary invasive lobular breast carcinoma
DAPI: PMCA2:γ-actin
CTS
CTS
G
Nuperphase tumor in primary prostate cancer
DAPI: PMCA2:γ-actin
AO
H
Primary breast cancer AMCC
DAPI: PMCA2:γ-actin
CCT
AO
I
Primary prostate AMCC with CCT degradation
DAPI: PMCA2:γ-actin
CCT
AO
CTS
J
Breast cancer with post-AMCC
DAPI: PMCA2:γ-actin
CCT
CTS
K
Breast cancer with CCT fragments
DAPI: PMCA2:γ-actin
CTS
CTS
L
Primary breast carcinoma with CCT degradation and oncocell apoptosis in CTs
DAPI: PMCA2:γ-actin
CT
CC
CT
CT
CTS
CT
CCT

Fig.S10

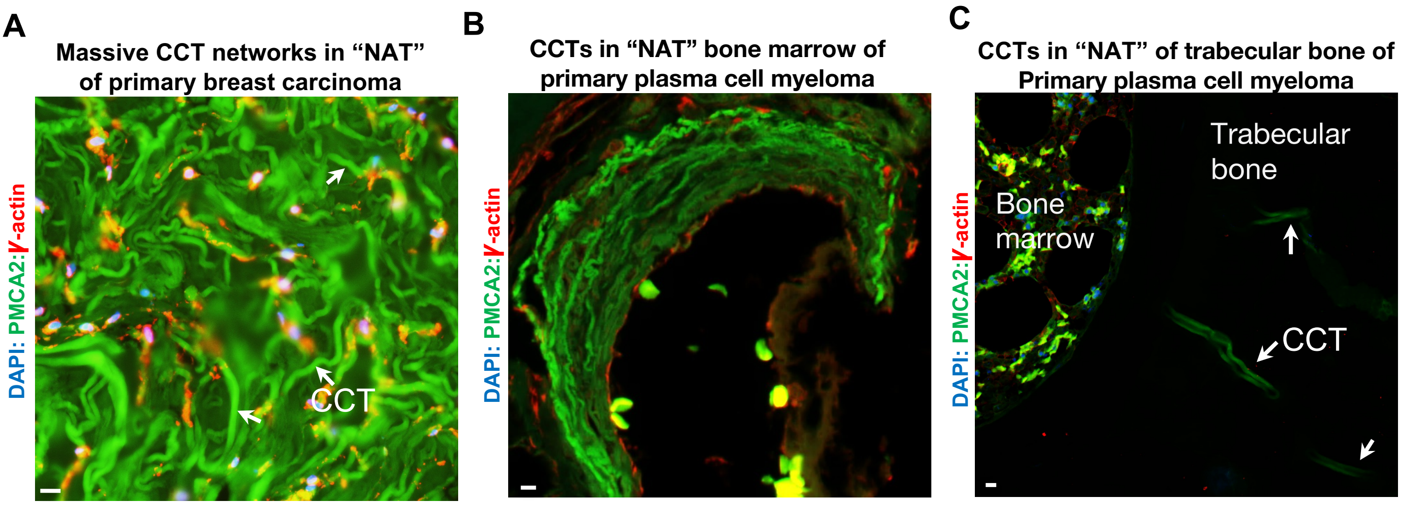


D

| Cancer types | Cancer subtypes | NAT tissue specimen number | Number of NAT tissues with CCT | % of paracancer tissue with CCT | CCT density/mm$^2$ |
|---|---|---|---|---|---|
| Bladder | 2 | 10 | 10 | 100 | 72-101 |
| Blood cancer in bone marrow | 2 | 20 (in trabecular bone) | 20 (in trabecular bone) | 100 | 2-10 |
| Blood cancer in bone marrow | 2 | 10 | 10 | 100 | 35-65 |
| Breast | 25 | 356 | 356 | 100 | 78-114 |
| Esophagus | 1 | 5 | 5 | 100 | 72-103 |
| Colon | 7 | 121 | 121 | 100 | 71-109 |
| Intestine | 1 | 3 | 3 | 100 | 78-107 |
| Kidney | 2 | 6 | 6 | 100 | 75-112 |
| Liver | 6 | 89 | 89 | 100 | 67-103 |
| Lung | 5 | 65 | 65 | 100 | 82-112 |
| Ovary | 2 | 12 | 12 | 100 | 79-114 |
| Pancreas | 6 | 96 | 96 | 100 | 81-107 |
| Stomach | 6 | 78 | 78 | 100 | 79-113 |
| Uterus | 1 | 5 | 5 | 100 | 81-102 |
| Total | 68 | 876 | 876 | 100 | |

Fig.S11

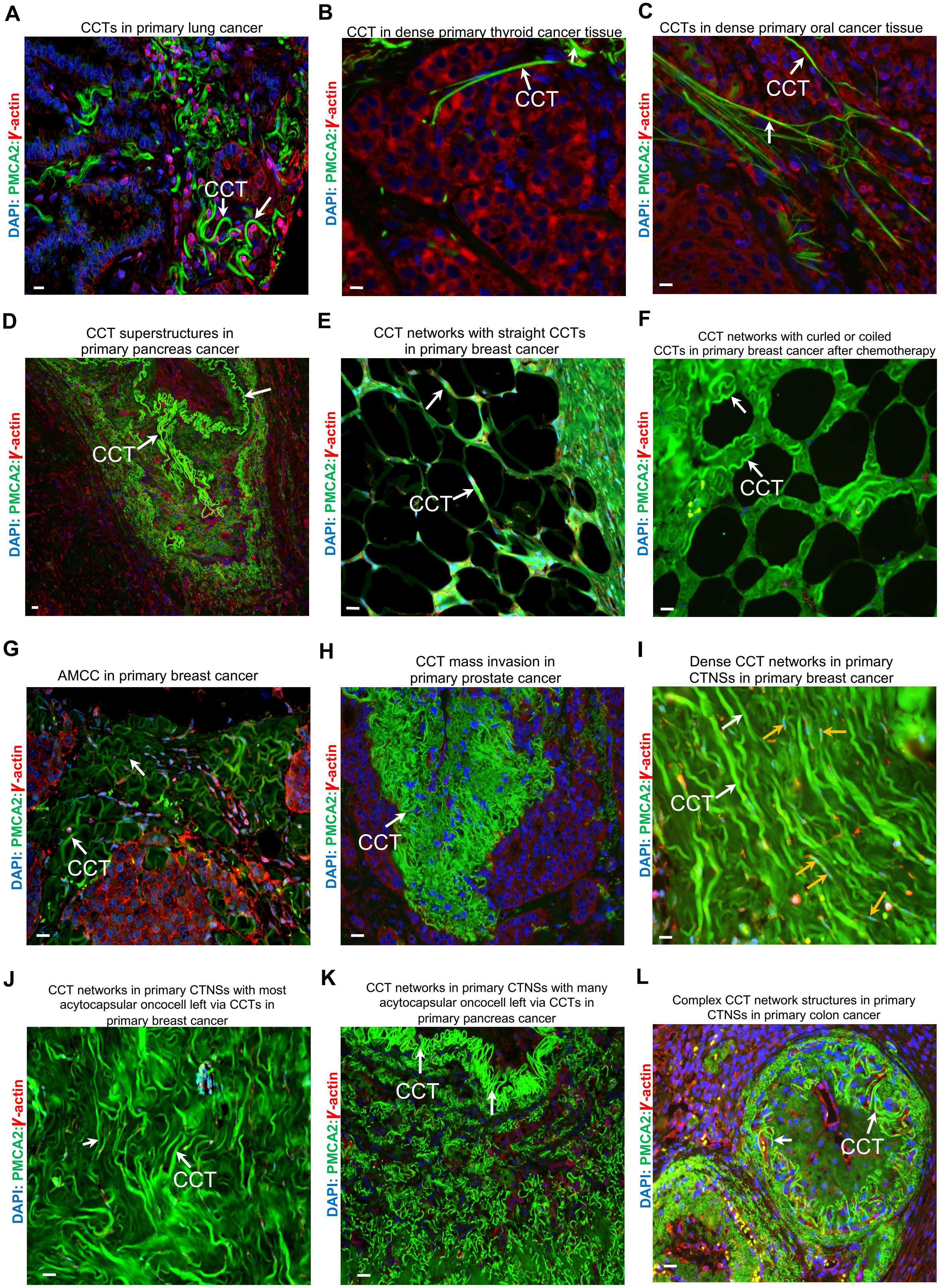
A
CCTs in primary lung cancer
DAPI: PMCA2:γ-actin
CCT
B
CCT in dense primary thyroid cancer tissue
DAPI: PMCA2:γ-actin
CCT
C
CCTs in dense primary oral cancer tissue
DAPI: PMCA2:γ-actin
CCT
D
CCT superstructures in primary pancreas cancer
DAPI: PMCA2:γ-actin
CCT
E
CCT networks with straight CCTs in primary breast cancer
DAPI: PMCA2:γ-actin
CCT
F
CCT networks with curled or coiled CCTs in primary breast cancer after chemotherapy
DAPI: PMCA2:γ-actin
CCT
G
AMCC in primary breast cancer
DAPI: PMCA2:γ-actin
CCT
H
CCT mass invasion in primary prostate cancer
DAPI: PMCA2:γ-actin
CCT
I
Dense CCT networks in primary CTNSs in primary breast cancer
DAPI: PMCA2:γ-actin
CCT
J
CCT networks in primary CTNSs with most acytocapsular oncocell left via CCTs in primary breast cancer
DAPI: PMCA2:γ-actin
CCT
K
CCT networks in primary CTNSs with many acytocapsular oncocell left via CCTs in primary pancreas cancer
DAPI: PMCA2:γ-actin
CCT
L
Complex CCT network structures in primary CTNSs in primary colon cancer
DAPI: PMCA2:γ-actin
CCT

**Fig.S12**

**A**

Cytocapsular oncocell metastasis via CCTs in primary prostate cancer

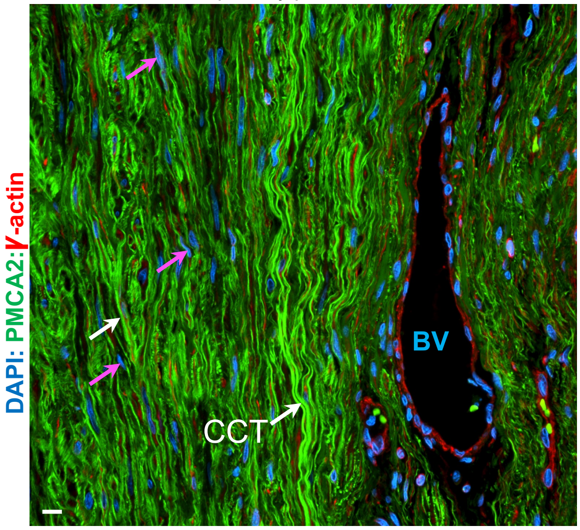


**B**

Comparison of cytocapsular tubes and humoral vessels in primary cancer niche, normal tissue adjacent to the tumor (NAT), and secondary cancer niche.

| Average ratios of the density of cytocapsular tubes to that of humoral vessels (blood vessels plus lymph vessels ) | | | |
|---|---|---|---|
| Cancer types | Primary cancer niche | NAT | Secondary cancer niche |
| Breast | 226 ± 4 | 277 ± 7 | 232 ± 7 |
| Colon | 212 ± 5 | 234 ± 6 | 256 ± 5 |
| Lung | 204 ± 7 | 238 ± 6 | 153 ± 7 |
| Pancreas | 289 ± 6 | 205 ± 3 | 261 ± 7 |
| Prostate | 223 ± 5 | 242 ± 5 | 136 ± 9 |
| Stomach | 205 ± 4 | 265 ± 8 | 236 ± 7 |

**C**

CCT invasion into micro blood vessels and release incytocapsular oncocell (ICO) into blood in primary prostate cancer

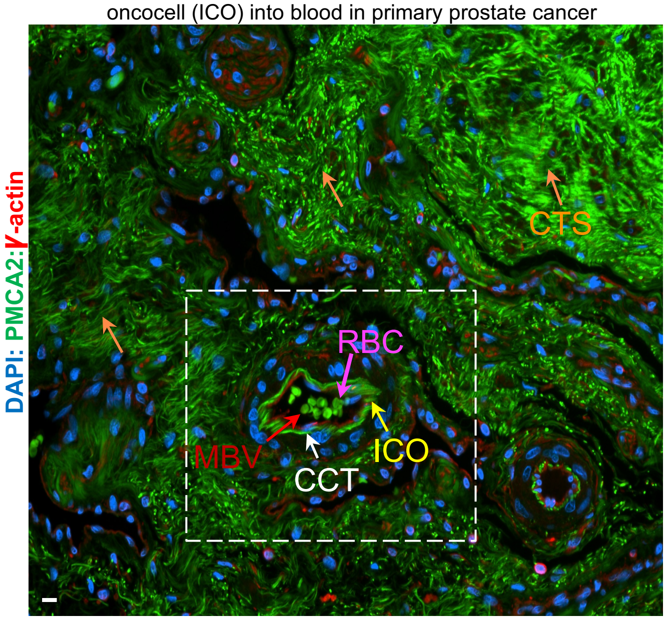


**D**

Enlarged image of CCT invasion into micro blood vessels and incytocapsular oncocell (ICO) in releasing into blood vessels

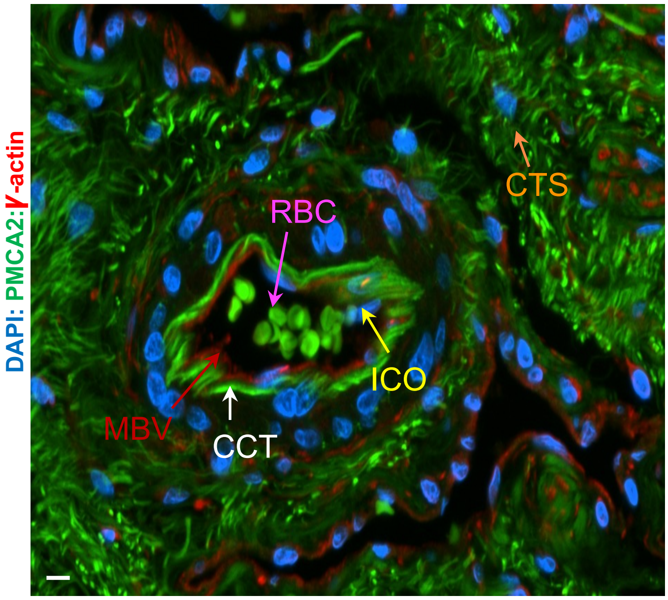


**E**

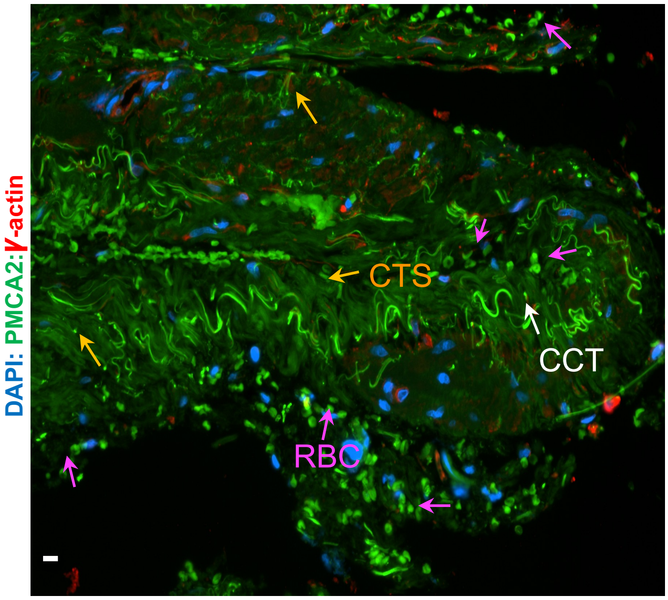


**F**

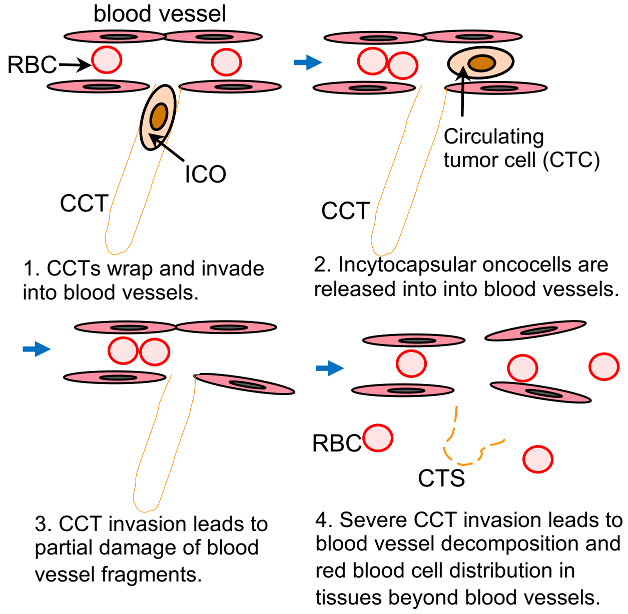

Fig.S13

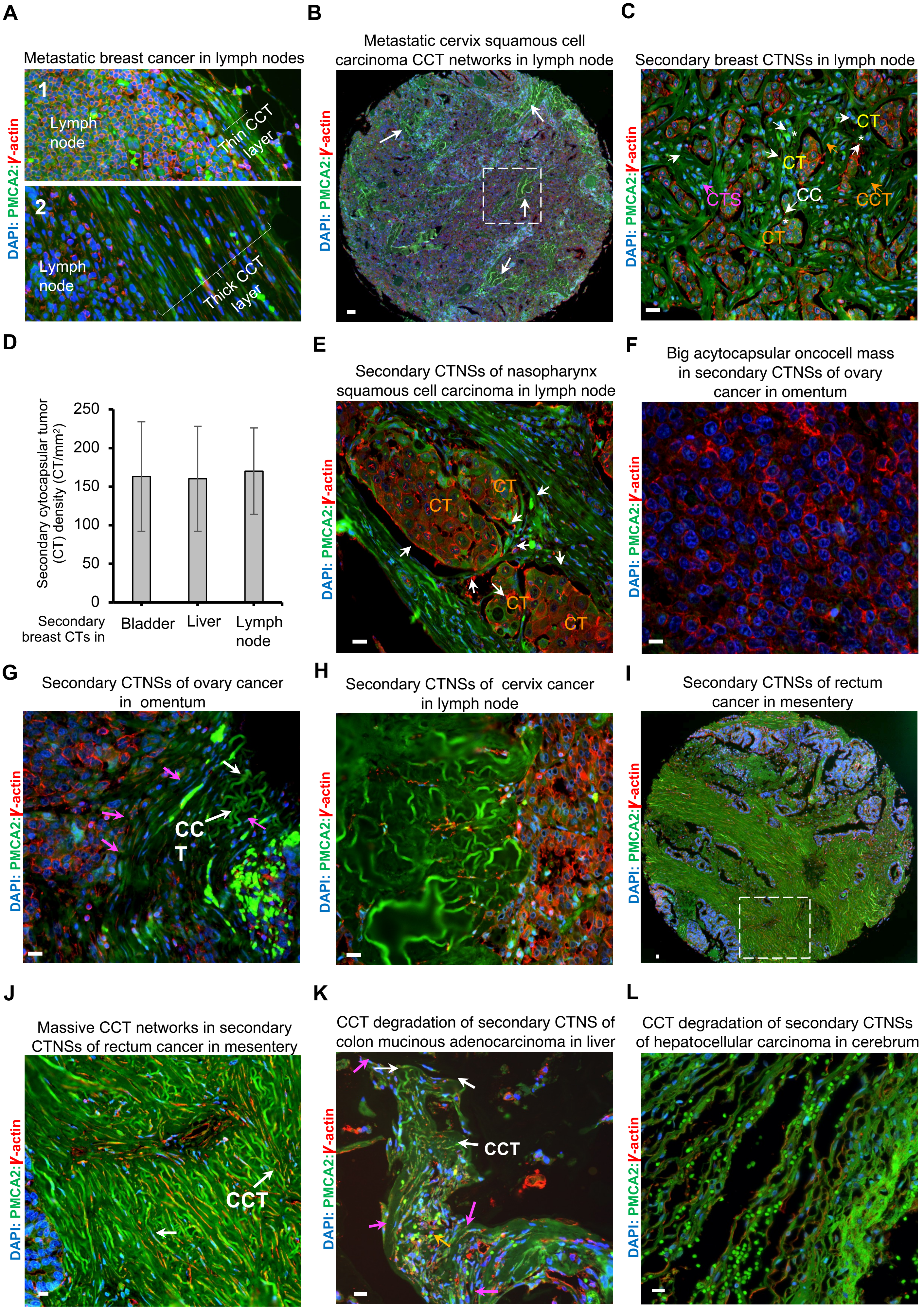
A
Metastatic breast cancer in lymph nodes
1
Lymph node
Thin CCT layer
2
Lymph node
Thick CCT layer
DAPI: PMCA2:γ-actin
B
Metastatic cervix squamous cell carcinoma CCT networks in lymph node
DAPI: PMCA2:γ-actin
C
Secondary breast CTNSs in lymph node
CT
CT
CTS
CC
CCT
CT
DAPI: PMCA2:γ-actin
D
Secondary cytocapsular tumor (CT) density (CT/mm2)
250
200
150
100
50
0
Secondary breast CTs in
Bladder
Liver
Lymph node
E
Secondary CTNSs of nasopharynx squamous cell carcinoma in lymph node
CT
CT
CT
CT
DAPI: PMCA2:γ-actin
F
Big acytocapsular oncocell mass in secondary CTNSs of ovary cancer in omentum
DAPI: PMCA2:γ-actin
G
Secondary CTNSs of ovary cancer in omentum
CCT
DAPI: PMCA2:γ-actin
H
Secondary CTNSs of cervix cancer in lymph node
DAPI: PMCA2:γ-actin
I
Secondary CTNSs of rectum cancer in mesentery
DAPI: PMCA2:γ-actin
J
Massive CCT networks in secondary CTNSs of rectum cancer in mesentery
CCT
DAPI: PMCA2:γ-actin
K
CCT degradation of secondary CTNS of colon mucinous adenocarcinoma in liver
CCT
DAPI: PMCA2:γ-actin
L
CCT degradation of secondary CTNSs of hepatocellular carcinoma in cerebrum
DAPI: PMCA2:γ-actin

**Fig. S14**

1.Normal cell

i ↓ Transformation

2.Acytocapsular oncocell (mass)

ii ↓ Devolution

3.Cytocapsular oncocell

iii ↓ Proliferation

4. Primary cytocapsular tumors

iv ↓ CCT generation, elongation, and branching morphogenesis

5. Primary CCT networks

v ↓ CTNS formation in primary niche

6.Primary CTNSs

vi ↓ Metastasis via CCT networks

7. Secondary cytocapsular tumors

vii ↓ CCT branching morphogenesis and new CT growth

8. Secondary CCT networks

viii ↓ CTNS formation in secondary niches

9. Secondary CTNS

ix ↓ Primary and secondary CTNS integration

10.Integrated primary and secondary CTNSs

x ↓ Cytocapsular oncocell secondary metastasis, new CT growth, and CTNS expansion; CC/CCT degradation and regeneration; AMCC formation, oncocell ecellulation and alloentry; therapy caused tumor disperse and/or CTNS fragment damages followed by new CT growth, CTNS recovering and expansion;

11. Dynamic integrated CTNSs

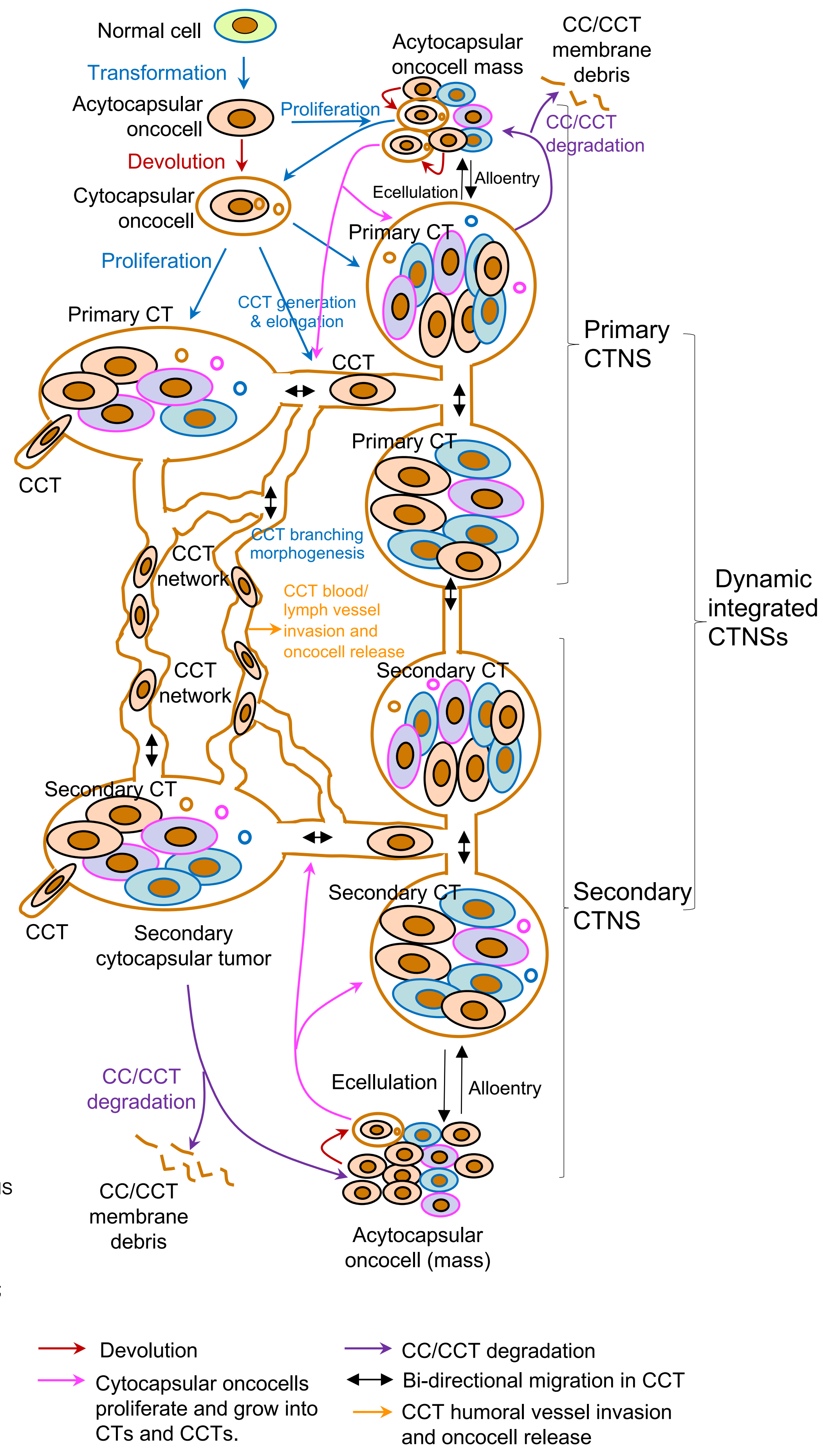
